# Excess prenatal folic acid supplementation alters cortical gene expression networks and electrophysiology

**DOI:** 10.1101/2025.05.07.652681

**Authors:** Viktoria Haghani, Sara Mohsen Ali, Noemi Cannizzaro, Mandar M. Patil, Paula DM. Sullivan, Ammara Rehamn, Ralph Green, Roy Ben-Shalom, Janine M. LaSalle, Konstantinos Zarbalis

**Author notes:** Corresponding authors, email, phone number.

## Abstract

Folate is crucial for various biological processes, with deficiencies during pregnancy being linked to increased risk for neural tube defects and neurodevelopmental disorders. As a proactive measure, folic acid fortification in foods has been mandated in many countries, in addition to dietary supplementation recommendations during pregnancy. However, the risks of excess prenatal folic acid supply have yet to be fully understood. To better appreciate *in utero* molecular changes in mouse brain exposed to 5-fold folic acid excess over normal supplementation, we investigated the transcriptome and methylome for alterations in gene networks. RNA-seq analysis of cerebral cortex collected at birth, revealed significant expression differences in 646 genes with major roles in protein translation. Whole genome bisulfite sequencing revealed 910 significantly differentially methylated regions with functions enriched in glutamatergic synapse and glutathione pathways. To explore the physiological consequences of excess prenatal folic acid exposure, we applied high-density microelectrode arrays to record network-level firing patterns of dissociated cortical neurons. Folic acid excess-derived cortical neurons exhibited significantly altered network activity, characterized by reduced burst amplitude and increased burst frequency, indicating compromised network synchronization.

These functional deficits align with the observed molecular alterations in glutamatergic synapse pathways, underscoring the potential for excess prenatal folic acid exposure to disrupt developing metabolic and neurological pathways.

## Introduction

Folate (vitamin B_9_) is an essential vitamin found in many foods, including dark leafy greens, fruits, legumes, seafood, meat, and more^1,2^. Folate is required for numerous biological processes, such as red blood cell production^3^, formation of the neural tube during fetal development^4–8^, DNA synthesis and repair^3,9,10^, and DNA methylation^10,11^. It has also been shown to have a protective effect against neurodevelopmental disorders, such as autism spectrum disorders (ASD) when taken in the first month of pregnancy, associated with maternal exposure to biological and chemical contaminants^12–19^. Due to its vital role in these processes, folate deficiency has been linked to several adverse health conditions, including an increased risk of miscarriages^4,20^, neural tube defects^3^, and increased risk of ASD^21–23^.

The neural tube, the precursor to the central nervous system, forms in humans during the first 28 days of development^24^, before many females become aware of their pregnancy^25^. To address this concern and ensure that pregnant females receive sufficient folate to support neural tube development among other developmental processes, folic acid (FA) fortification became a standard public health measure in many countries, including the United States^26^ and Canada^27^ in 1998, Australia and New Zealand in 2009^28^, South Africa in 2003^29^, Chile in 2000^30,31^, Argentina in 2003^32^, Brazil in 2004^33^, and many more. Consequently, a notable reduction in neural tube defects and birth defects was documented^34^, with one study demonstrating a 72% protective effect against neural tube defects^35^. In addition to FA fortification typically added to grains, it is common practice for pregnant females to take prenatal vitamins that further supplement the diet with FA^36^.

While the benefits of FA supplementation during neurodevelopment are well documented, emerging evidence suggests that FA excess (FAE) may have unintended negative consequences^37^, such as disruptions to the immune system^38^, alterations in DNA methylation patterns^39,40^, metabolic perturbations^41–45^, and increased risk of certain cancers^46–48^. Additionally, there are indications that FAE could interfere with the regulation of homocysteine levels^49,50^, which is associated with cardiovascular disease^51^.

Furthermore, some epidemiological studies have also postulated risks associated with high prenatal FA taken at any time in pregnancy to neurodevelopment and neurodevelopmental disorders^52–56^. Of particular interest, data from the Boston Birth Cohort showed a’U shaped’ relationship between maternal multivitamin supplementation frequency and ASD likelihood. A positive association between maternal plasma folate levels at birth and autism likelihood was identified^57^ mostly implicating high plasma concentrations of unmetabolized FA^58^.

A conceivable mechanism by which FAE may adversely impact neuronal function is its influence on neuronal circuit development. The formation of such circuits involves timely structured synaptogenesis and pruning to shape both local and long-range connectivity patterns^59^. During normal brain development, these mechanisms facilitate the maturation of neural networks through selective strengthening and weakening of synaptic connections^60^. Disruptions in these fundamental developmental processes have been implicated in various neurodevelopmental disorders, particularly ASD, where altered patterns of neural connectivity have been observed^61^. These potential risks highlight the need to investigate the consequences of FAE.

In this study, we evaluated the effects of 5-fold FAE on gene pathways and networks in the neonatal mouse brain, with particular emphasis on changes in gene expression and DNA methylation that could contribute to neurodevelopmental impairments. By comparing RNA-seq and whole genome bisulfite sequencing (WGBS) data between the control and FAE groups, we uncovered dysregulated genes and epigenetic alterations to translational regulation, glutathione metabolism, and glutamatergic neuronal synapses, all functions relevant to neurodevelopment. To further test possible functional consequences of identified molecular dysregulations, we applied high-density microelectrode array (HD-MEA) technology to enable the detailed examination of neuronal network firing patterns. Our results confirm diminished burst amplitudes and synchronicity of neuronal networks formed by FAE-derived dissociated cells obtained from newborn pups.

## Materials & Methods

### Animal Husbandry and Diets

Mice were kept in facilities accredited by the Association for Assessment and Accreditation of Laboratory Animal Care International. All procedures were conducted following protocols approved by the Institutional Animal Care and Use Committee at the University of California, Davis, ensuring compliance with ethical guidelines for animal research. Dietary groups were established by feeding C57BL/6NJ dams Clifford/Koury-based L-amino acid-defined rodent diets^62^ (Dyets Inc., Bethlehem, PA) with specified folic acid levels, starting two weeks before mating and continuing through pregnancy. The breeding pairs remained on their assigned folate-controlled diets until the pups were collected. The experimental groups included: (1) a control group receiving 2 mg/kg of folic acid and (2) a high-folate group receiving 10 mg/kg (equivalent to 5 times the control amount). The control diet with 2 mg folic acid/kg chow satisfies the experimentally defined daily folate requirement for rodents^63^, as recommended by the American Institute of Nutrition^64^, corresponding to the human recommended intake of 400 μg/day.

### Tissue Collection and Extraction

Newborn pups (P0-P2) were collected within hours after birth. Each pup was decapitated, and the cerebral cortex was rapidly dissected, placed on ice in tubes with brain preservation media which consisted of Hibernate-A and 2% 1x B-27 Supplement, and promptly stored at 4°C until further processing. Genomic DNA and total RNA were simultaneously extracted from the same cortical tissue sample using the Qiagen AllPrep DNA/RNA/miRNA Universal Kit, following the manufacturer’s protocol to maximize yield. The DNA and RNA were separately eluted and assessed for quality and concentration using a Nanodrop spectrophotometer, with additional verification on agarose gels. DNA and total RNA were submitted to Novogene Corporation Inc. (Sacramento, CA) for RNA-seq and WGBS sequencing. PCR library selection was performed and paired-end sequencing was conducted on the Illumina NovaSeq X Plus platform. Sequencing generated paired-end reads of 150 bp in length, targeting approximately 30-40 million reads per sample for RNA-Seq and 40-50 million reads per sample for WGBS. Libraries were sequenced to a depth of >20X coverage for WGBS. Raw sequence data output was generated in FASTQ format, with quality scores provided in Phred format (Qphred =-10log10(e), where’e’ represents the sequencing error rate.

Sequencing results of 7 cerebral cortical samples from the control group and 9 samples from the FAE group were analyzed for this study.

### RNA-seq Bioinformatics Analysis

Raw sequence data was trimmed using Trim Galore^65^ (v0.6.10) and aligned to the mmEnsemble107 genome using STAR^66^ (v2.5.2b) with the Mus_musculus_Ensemble_107_new GTF file for annotation. The trimmed sequence data was aligned to the hg19, hg38, Lambda, mm10, PhiX, rheMac10, and rn6 genomes using FastQ Screen^67^ (v0.15.3) to assess sample origins and screen for possible contamination. Samtools^68^ (v1.11) was used to index the aligned BAM files. The proportion of reads mapping to the X vs. Y chromosome was assessed to determine the sex of the samples. MultiQC^69^ (v1.14) was run for all raw and processed data to evaluate data integrity and overall quality.

To identify differentially expressed genes (DEGs), raw count data output by STAR was read into a Jupyter Notebook^70^ (v7.0.7) running R^71^ (v4.3.2) via IRkernel^72^ (v1.3.2). Two DEG analyses were conducted. The first analysis contrasted control (n = 7) vs. FAE (n = 9) with sex as a covariate. The second analysis was sex-segregated, containing control females (n = 4), FAE females (n = 6), control males (n = 3), and FAE males (n = 3). The following analyses were conducted: control female vs. FAE female, control male vs. FAE male, FAE female vs. FAE male, and control female vs. control male. The raw count data were normalized in R. The data were filtered based on criteria described by Chen et al. 2016^73^. Briefly, genes were only kept if they had a minimum count-per-million in at least 70% of samples from the smallest group size and a minimum total count across all samples. The normalized and filtered data were log-transformed and subjected to voom transformation, then fitted to a linear model using limma^74^ (3.58.1) in conjunction with edgeR^75^ (v4.0.16). Contrasts were defined for each pairwise comparison following the convention of FAE minus control where relevant for consistent interpretation of log fold changes, and the estimated contrasts for each gene were calculated.

Finally, empirical Bayes moderation was applied, followed by multiple testing adjustment using the Benjamini-Hochberg method to identify significant DEGs.

In addition to DEG analysis, weighted gene correlation network analysis (WGCNA) was also conducted using the WGCNA package^76^ (v1.72.5) to identify co-expression modules of genes, relate these modules to external traits, and identify key hub genes that may play significant roles in the biological processes associated with FAE. Normalized count data were used to generate WGCNA in R. Soft power thresholding was performed, and a power of 17 was selected because the SFT.R.sq value was 0.80600, indicating a strong scale-free topology for the network. This choice of power ensures that the resulting network accurately reflects the underlying biological relationships among the genes. Blockwise modules were generated using a power of 17 to construct the network, with parameters set to optimize module detection for genes exhibiting coordinated expression patterns. Module eigengenes were calculated for modules, representing the combined expression patterns of the genes within each module.

Then, module membership was assessed by calculating the biweight midvariance correlation and associated p-values between the genes and the module eigengenes, enabling the evaluation of each gene’s association with the identified modules. Pearson correlation coefficients and associated p-values were calculated between the module eigengenes and the sample metadata, facilitating the assessment of the relationships between gene modules and external traits, namely: condition only (control vs. FAE), sex only, control female vs. FAE female, control male vs. FAE male, and FAE female vs. FAE male.

Gene ontology (GO) was carried out for both DEG and WGCNA results. Biomart^77^ (v2.58.2) was used to assign Entrez gene IDs to Ensembl genes. For the DEG GO analysis, genes were filtered to adjusted p-values < 0.05 and submitted to the following Enrichr^78^ (v3.2) databases: GO Biological Process 2023, GO Cellular Component 2023, GO Molecular Function 2023, KEGG 2019 Mouse, Panther 2016, Reactome 2016, and RNA-seq Disease Gene and Drug Signatures from GEO. Significant DEGs were categorized as upregulated or downregulated with FAE exposure based on the directionality of log-fold changes. Plots were created using a combination of ggplot2^79^ (v3.4.4) and Enrichr to depict the top enriched terms from the Enrichr databases for all significant DEGs, as well as for upregulated and downregulated DEGs. For WGCNA, genes within each module were submitted as lists to the same Enrichr databases used for the DEG GO analysis. Enriched terms were plotted using ggplot2 and Enrichr.

### WGBS Bioinformatics Analysis

Raw sequence data was processed using Epigenerator^65,69,80–82^. Briefly, sequence data from multiple lanes for the same sample were merged. The data were trimmed using Trim Galore^65^ (v0.6.10). The trimmed sequence data were aligned to the hg19, hg38, Lambda, mm10, PhiX, rheMac10, and rn6 genomes using FastQ Screen^67^ (v0.15.3) to assess sample origins and screen for possible contamination. Trimmed data were aligned to the mm10 genome and deduplicated using Bismark^81^ (v0.24.0). Data were sorted by chromosome coordinate and insert size metrics were collected using Picard^83^ (v2.27.5). Bismark was used to calculate nucleotide coverage and compare it to average genome composition, determine methylation status for genomic cytosines, and generate cytosine reports, detailing cytosine coverage and methylation status. MultiQC^69^ (v1.14) was run to ensure sequence quality for processed data. Five samples were excluded from analysis due to low alignment rates (0.1%), which was caused by improper insert sizes (i.e. paired-end fragments are either too short or too long, resulting in insufficient sequence content). The remaining samples were comprised of the following: control females (n = 3), control males (n = 4), FAE females (n = 2), and FAE males (n = 3).

Cytosine reports were used as inputs for DMRichR^84–86^ (v1.7.1), which identifies differentially methylated regions (DMRs) and performs enrichment analysis. The analysis was conducted as control vs. FAE with sex as a covariate in the analysis. DMRs were identified when at least 50% of samples in a group exhibited coverage in a specified region displaying at least 10% difference in methylation levels between groups. Additionally, cytosine reports were used as inputs for Comethyl^87^ (v1.3.0), which performs WGCNA to identify modules of genomic regions with correlated methylation patterns. A minimum coverage of 2 was required for a CpG site to be retained in the analysis with at least 70% of samples meeting the coverage requirement.

Regions, which require a minimum of 3 CpG sites within 150 bp, were filtered to retain those with a minimum coverage of 8 across samples. Modules were generated using a mergeCutHeight of 0.4 with a minimum module size of 60, meaning that clusters of genes were combined if their dissimilarity was less than 0.4, and each module contains at least 60 genes.

This allowed for the exploration of biologically relevant relationships between methylated regions, essentially identifying DMRs that may act together as a network due to the interconnectedness of genomic methylation changes.

### Integrative RNA-seq and WGBS Bioinformatics Analysis

Because the RNA-seq and WGBS data were derived from paired samples obtained from the same mice, the RNA-seq and WGBS data can be correlated. Correlations were conducted between RNA-seq data (transcripts per million, TPM) and WGBS data (percent methylation). To calculate RNA-seq TPM, we utilized normalized count data and the mm10 reference genome annotation (mm10.refGene.gtf). We summed the lengths of exons for each gene to obtain the total gene length, which was then converted to kilobases by dividing by 1,000. Reads per kilobase (RPK) were computed by dividing the normalized read counts by the gene lengths in kilobases. Finally, TPM was derived from the RPK values by normalizing each gene’s RPK to the total RPK across samples and multiplying by 1,000,000 (i.e. TPM = (RPK / Total RPK) * 1,000,000). To determine the WGBS percent methylation, a total count of unmethylated and methylated cytosines per gene were generated from the cytosine reports. Percent methylation was calculated as the number of methylated cytosines divided by the total cytosines (methylated and unmethylated) times multiplied by 100 (i.e. percent methylation = (methylated cystosines / (methylated cytosines + unmethylated cytosines)) * 100).

To assess the correlation between gene expression and DNA methylation, samples were grouped into control and FAE groups, with each sample in the corresponding group being treated as an observation containing paired data (TPM and percent methylation). Spearman correlation coefficients and their p-values were calculated for every gene in the data sets. Then, overlapping significant DEGs and significant DMRs were identified and subject to a Fisher’s Exact Test to assess the significance of overlap.

### Cell Culture and Microelectrode Array Analysis

Primary cortical neuron-glia cultures were established from P0-P2 newborn mouse brain tissue from one dam per dietary group (control and FAE). Following decapitation under approved anesthesia protocols, cortical tissue was enzymatically dissociated using papain (20 units/ml) and DNase-1 from bovine pancreas in Hibernate-A medium, followed by mechanical trituration to obtain single-cell suspensions.

Dissociated cells were plated at a density of 125,000 cells per chip on Maxwell Biosystems high-density microelectrode arrays (HD-MEAs) pre-coated with 0.07% polyethyleneimine and laminin (both Sigma-Aldrich, St. Louis, MO). Cultures were maintained in Neurobasal media supplemented with B27 and 10% horse serum for the first 24 hours, then switched to DMEM containing GlutaMax, D-glucose, sodium pyruvate, and 10% horse serum, with 50% media changes performed every three days.

Electrophysiological recordings were conducted between days *in vitro* (DIV) 7 and DIV 33, with recordings performed twice weekly to track development across three phases: early network formation (DIV 7-14), initial network maturation (DIV 14-21), and network stabilization (DIV 21-28). Recording sessions used temperature-equilibrated units maintained at 37°C with 5% CO_₂_ following a 10-minute equilibration period to ensure stable activity.

Each recording session began with an Activity Scan Assay using MaxLab Live software, which measured neuronal activity across all 26,400 electrodes for 30 seconds each.

Subsequently, the Network Assay module recorded simultaneous activity from 1,024 high-firing electrodes selected based on signal amplitude and spatial distribution for 300 seconds at 20 kHz sampling rate.

Data analysis was performed using custom MATLAB R2024 scripts and the MaxWell Biosystems MATLAB toolbox (v23.2). Raw voltage traces were bandpass filtered (300 - 3000 Hz) and thresholded at 5× the standard deviation of baseline noise for initial spike detection (for network-level analyses only). The detected spike rasters were converted into network activity by binning spike events and applying Gaussian smoothing to estimate the instantaneous population firing rate. Network parameters, including burst frequency and amplitude, were computed and statistically compared using unpaired two-tailed *t*-tests

## Results

### FAE leads to sex-specific transcriptional dysregulations associated with mitochondrial function and protein synthesis

For the sex combined analysis (control vs. FAE), 646 significant DEGs (adjusted p-value < 0.05) were identified (Supplementary Table 1). For the sex-stratified analyses, only the control female vs. FAE female analysis yielded significant DEGs (161, Supplementary Table 2), with the other comparisons (control male vs. FAE male, control male vs. control female) having no significant DEGs. Significant DEGs for the sex-adjusted combined and sex-stratified analyses were compared to identify overlapping genes (i.e. DEGs present in both analyses) that may be consistently affected by FAE (Figure 1, Supplementary Table 3). There were 267 upregulated DEGs unique to control vs. FAE, 228 downregulated genes unique to control vs. FAE, 10 downregulated DEGs overlapping between the two comparisons, 51 upregulated DEGs overlapping between the comparisons, 7 downregulated DEGs unique to control female vs. FAE female, and 3 upregulated DEGs unique to control female vs. FAE female. Because DEG contrasts were conducted as FAE minus control, “upregulated” refers to genes expressed more highly in the FAE group and “downregulated” refers to genes expressed lower in the FAE group compared to the control group.

**Figure 1.**
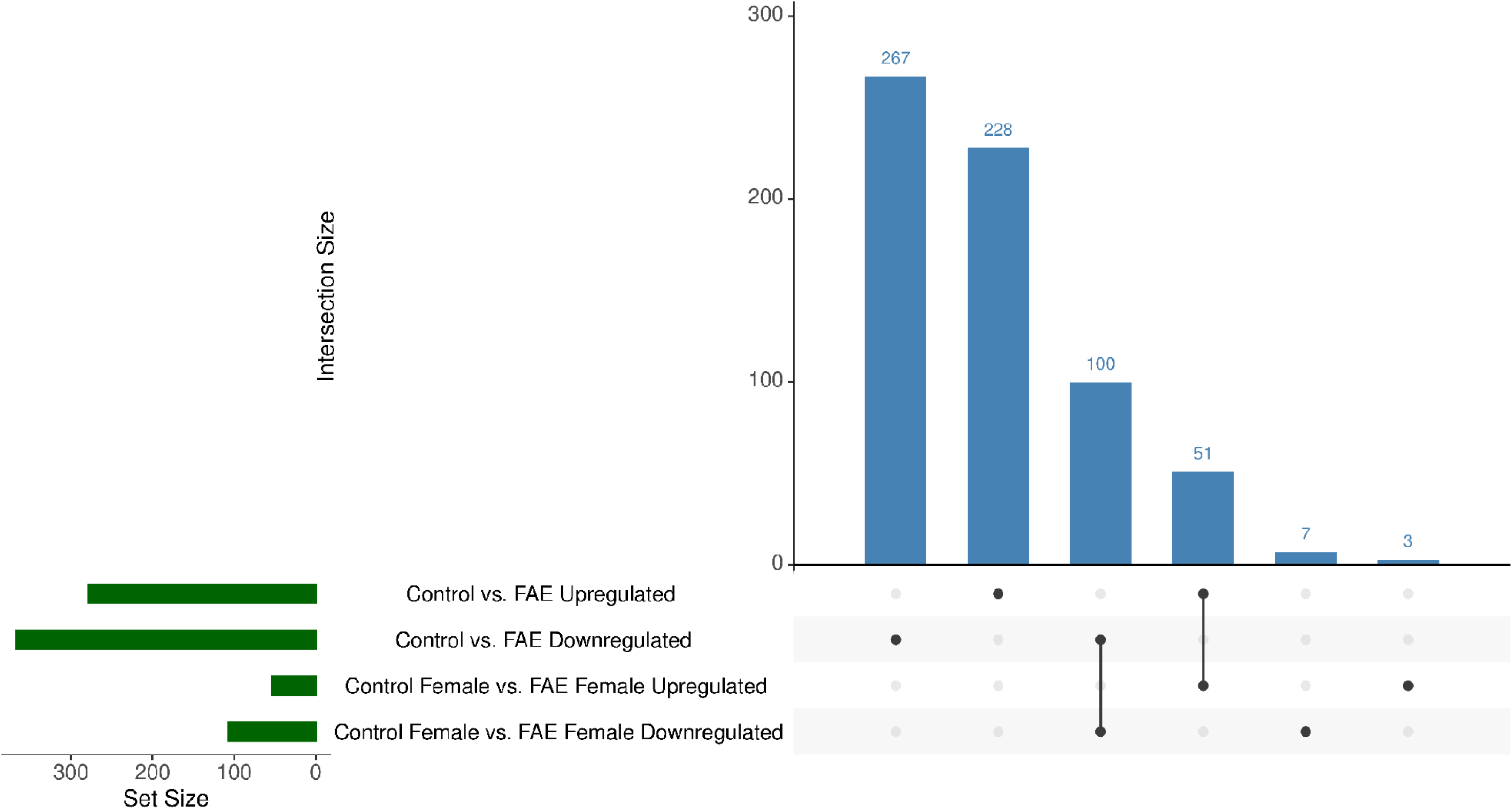
UpSet Plot of DEGs. Upregulated and downregulated DEGs were overlapped between the sex combined and sex-segregatedRNA-seq analyses. The number of upregulated and downregulated genes in each comparison was plotted, with specific emphasis on the intersections between the groups to discern sex-specific DEG results. This UpSet plot was made using R (v4.3.2) with an R kernel (r-irkernel v1.3.2) in a Jupyter Notebook (v7.0.7) with the following packages: UpSetR (v1.4.0), openxlsx (v4.2.5.2), readxl (v1.4.3), dplyr (v1.1.14), glue (v1.7.0), and ggplot2 (v3.4.4).

The 10 most significantly upregulated and downregulated DEGs were labelled for the control vs. FAE and control female vs. FAE female DEG analyses (Figure 2). A few overlaps were identified among these genes, including *Gm55094*, *Lnp1*, *Hhex*, and *Dcst2*, which are downregulated with prenatal FAE exposure. *Ackr2* overlaps for the top 10 upregulated genes. Notably, *Lnp1*, which is downregulated, is involved in lipid metabolism, and *Ackr2*, which is upregulated, is involved in the immune response.

**Figure 2.**
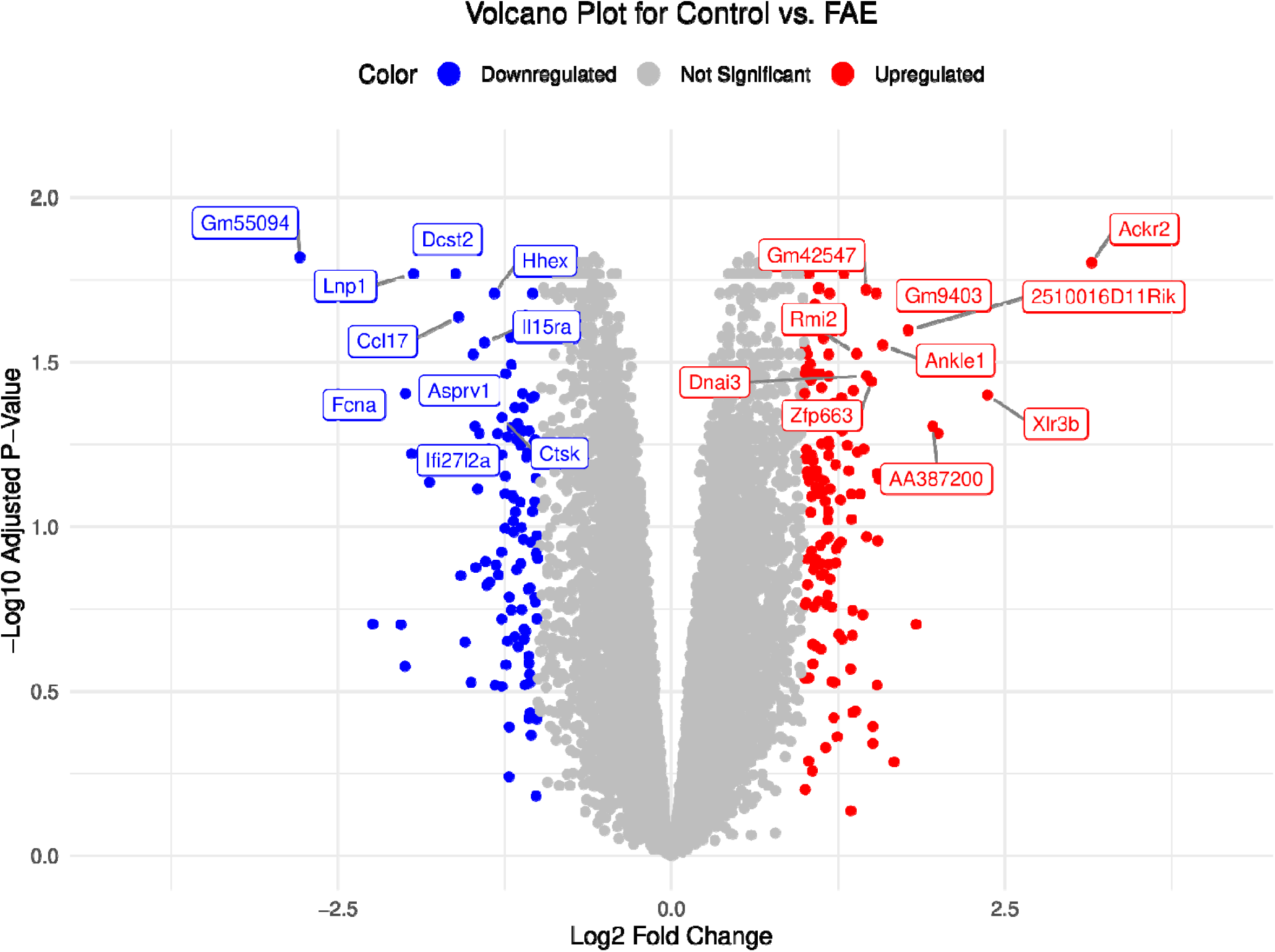
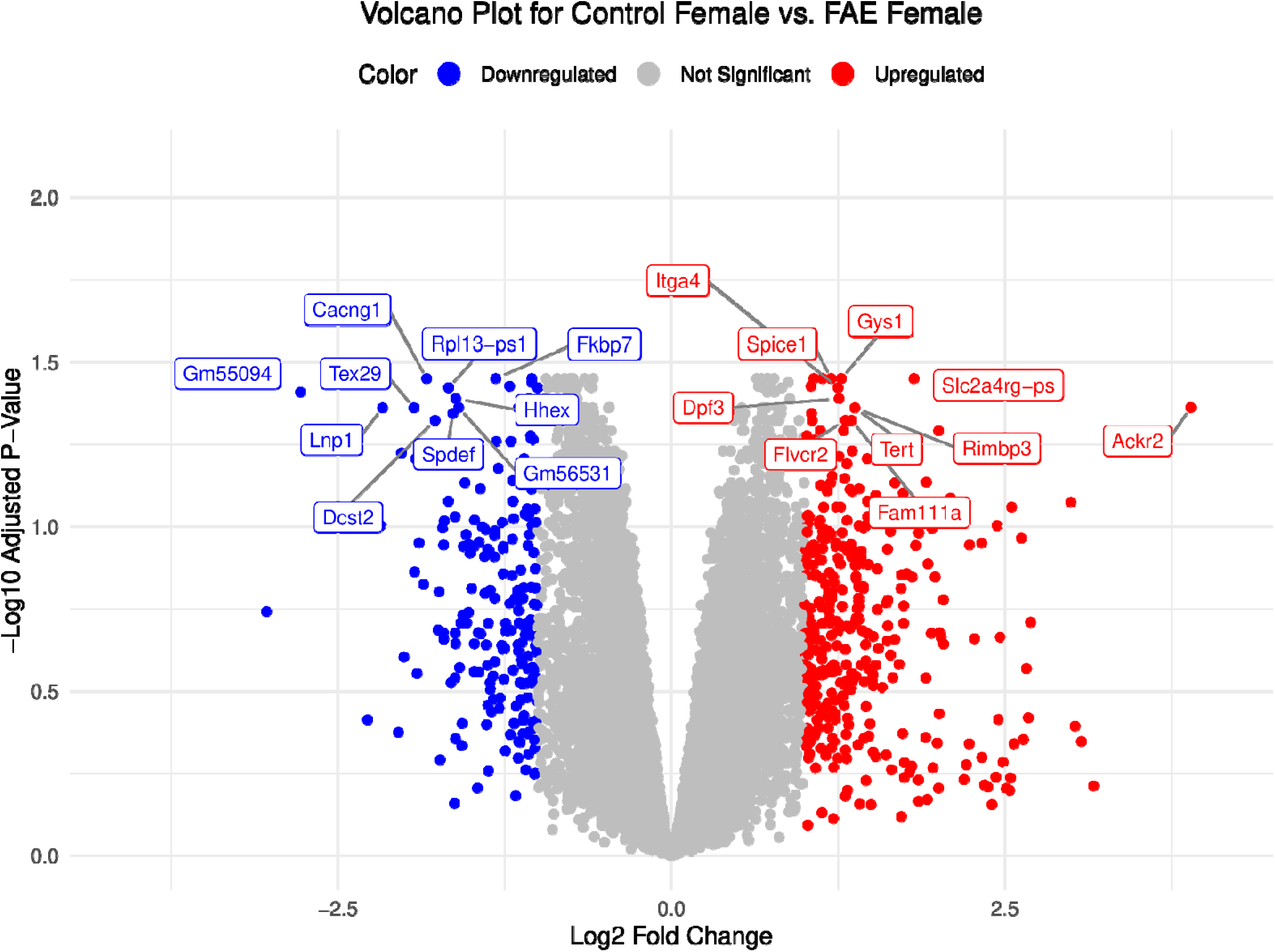
Volcano Plots of Top 10 Significant Upregulated and Downregulated DEGs. This figure displays volcano plots of the top 10 most significantly differentially expressed genes filtered for adjusted p-values less than 0.05 and a log fold change greater than 1 for **(A)** control vs. FAE and **(B)** control female vs. FAE female. The x-axis represents the log2 fold change in gene expression, while the y-axis indicates the-log10 of the adjusted p-values, allowing for the visualization of both the magnitude and significance of expression changes. Upregulated genes are marked in red, downregulated genes are shown in blue, and genes that are not significantly differentially expressed are colored gray. The top 10 upregulated and downregulated genes are also labeled by gene name. This volcano plot was made using R (v4.3.2) with an R kernel (r-irkernel v1.3.2) in a Jupyter Notebook (v7.0.7) with the following packages dplyr (v1.1.14), glue (v1.7.0), ggplot2 (v3.4.4), and ggrepel (v0.9.6).

Significant DEGs were used to conduct the GO analysis, which revealed several significant biological processes, cellular components, and molecular functions (Figure 3). Key biological processes corresponding to genes downregulated in FAE included cytoplasmic translation, proton motive force-driven mitochondrial ATP synthesis, translation, macromolecule and peptide biosynthetic processes, and oxidative phosphorylation, indicating a strong impact of FAE on protein metabolism. Notable cellular components identified were various mitochondrial structures, including the mitochondrial inner membrane and ribosomal complexes, suggesting alterations in mitochondrial function and protein synthesis. Additionally, molecular functions associated with NADH dehydrogenase activity, as well as binding to RNA, ribosomes, and various types of rRNA, were identified, indicating alterations in electron transport and ribosomal activity. GO terms corresponding to genes upregulated in FAE included sterol and cholesterol biosynthetic processes, indicating enhanced capacity for lipid metabolism and membrane synthesis. Additionally, positive regulation of phosphatidylinositol 3-Kinase signaling and DNA damage response suggest a potential role in cellular signaling and stress response pathways.

**Figure 3.**
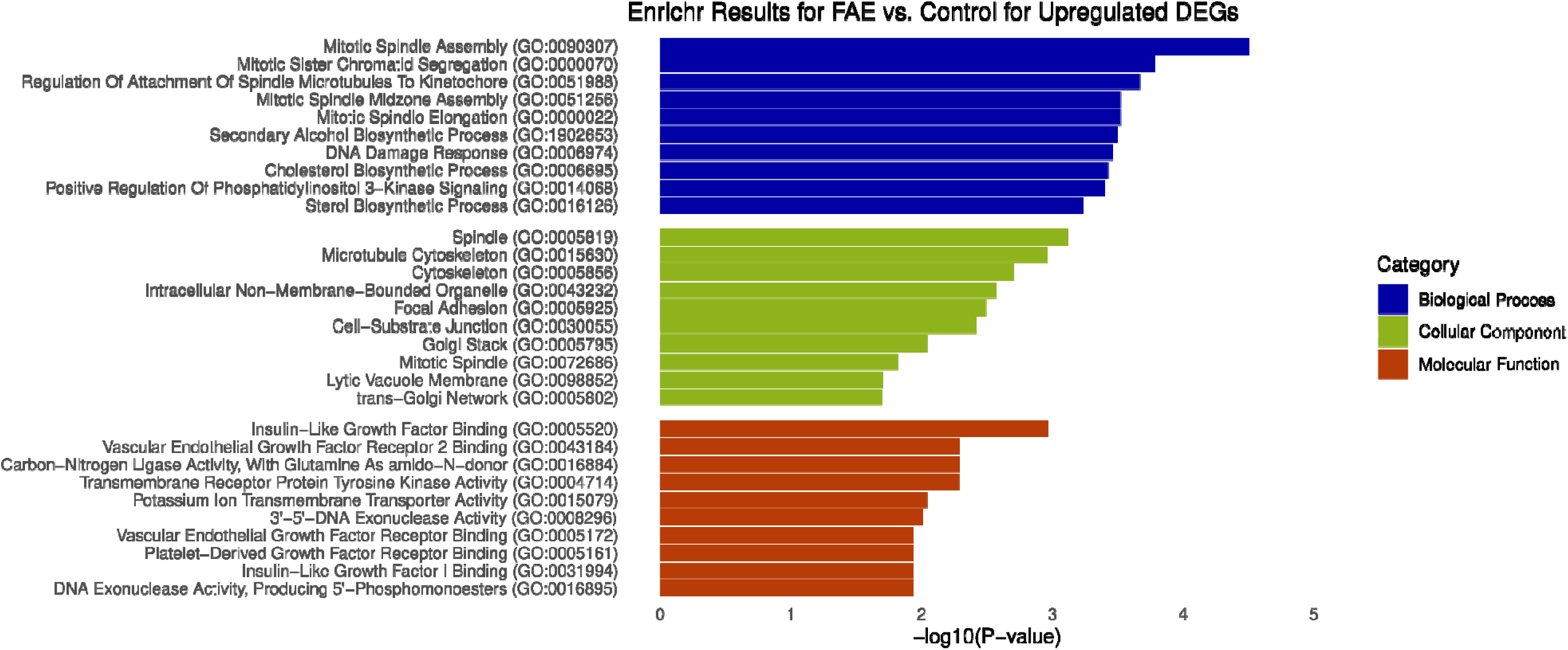
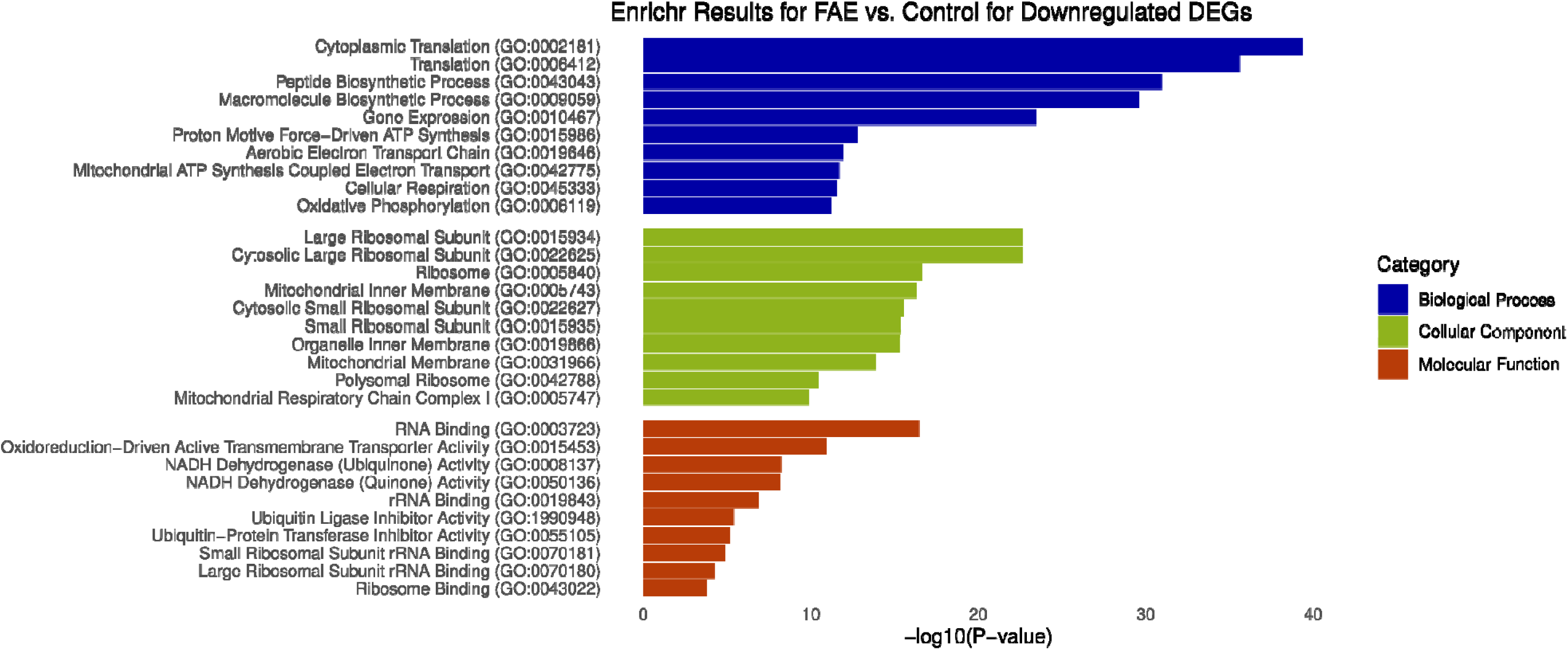
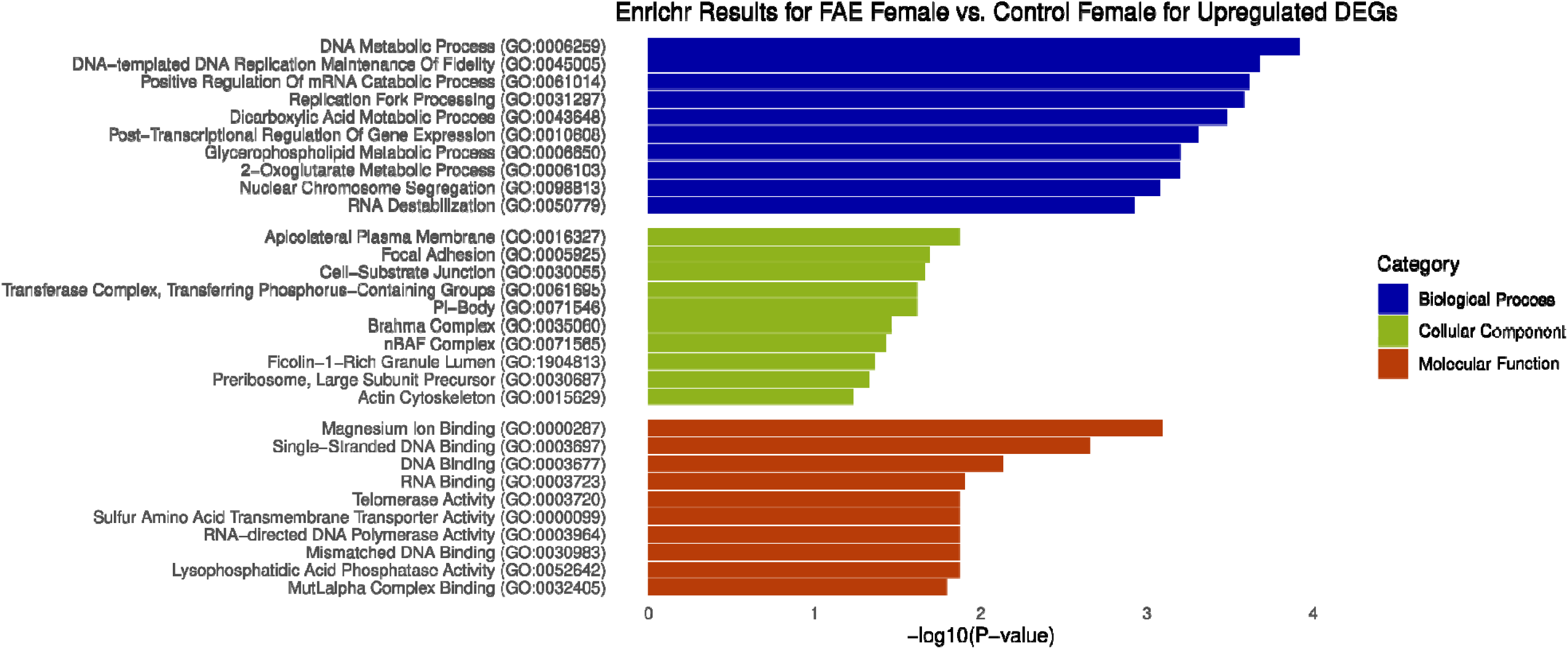
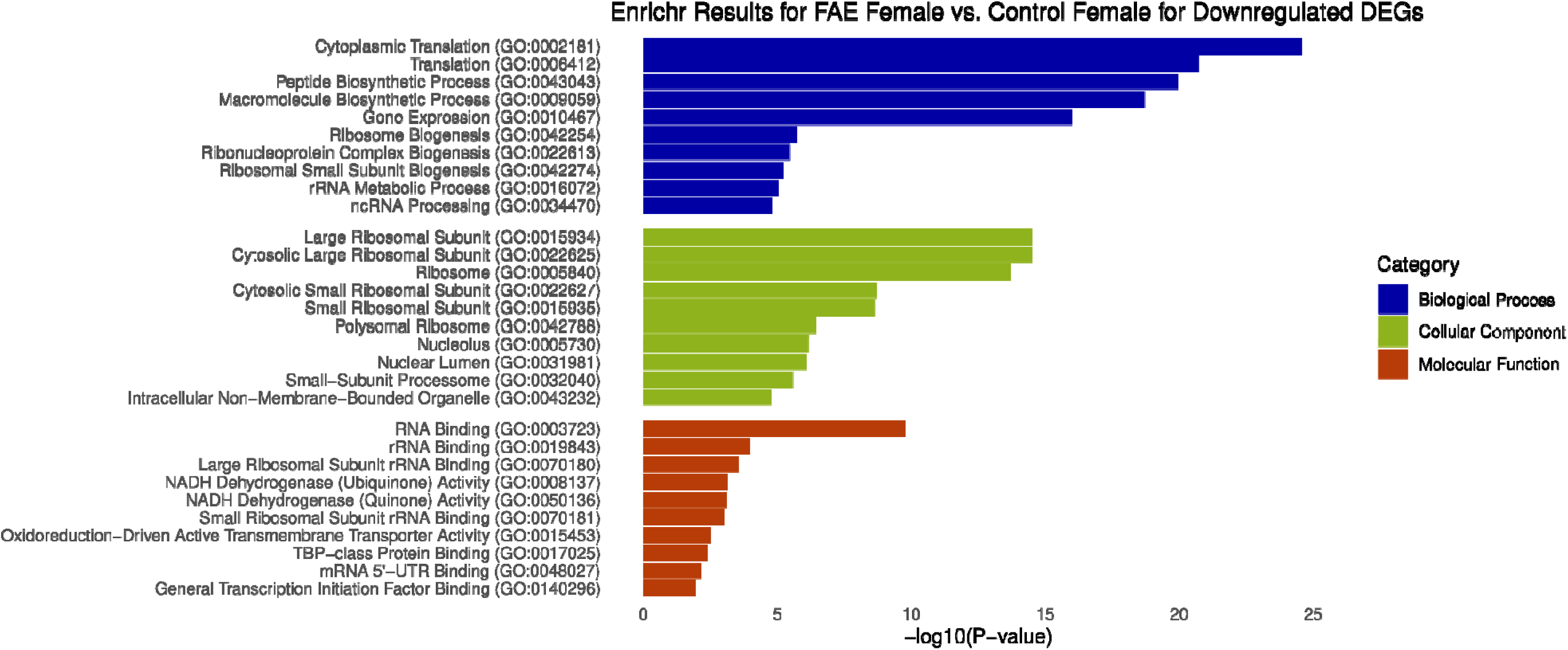
Enrichr Gene Ontology Analysis for DEGs. This figure displays the results of the gene ontology analysis for the top 10 enriched terms identified in the GO Biological Process 2023 (purple), GO Cellular Component 2023 (blue), and GO Molecular Function 2023 (green) Enrichr databases. **A** represents the results associated with upregulated genes for the control vs. FAE comparison, while **B** shows the results associated with downregulated genes for the control vs. FAE comparison. **C** represents the results associated with upregulated genes for the control female vs. FAE female comparison, while **B** shows the results associated with downregulated genes for the control female vs. FAE female comparison. The x-axis represents the-log10 of the p-values, indicating the significance of each term, with higher values reflecting greater significance. The y-axis lists the enriched terms. These bar plots were made using R (v4.3.2) with an R kernel (r-irkernel v1.3.2) in a Jupyter Notebook (v7.0.7) with the following packages: enrichr (v3.2), dplyr (v1.1.14), ggplot2 (v3.4.4), biomaRt (v2.58.2), openxlsx (v4.2.5.2), and readxl (v1.4.3).

Cellular components associated with genes upregulated in FAE include trans-Golgi network, lytic vacuole membrane, mitotic spindle, spindle, and cytoskeleton, which suggests a potential increase in cell growth and proliferation. Molecular functions for upregulated genes are involved in pathways such as DNA exonuclease activity, insulin-like growth factor binding, vascular endothelial growth factor receptor 2 binding, and 3’-5’ DNA exonuclease activity, which is also consistent with cell growth and proliferation.

The upregulated genes identified in the control female vs. FAE female analysis were associated with biological processes such as DNA metabolic process, DNA-templated DNA replication maintenance of fidelity, and replication fork processing, indicating major roles in DNA synthesis. Additionally, positive regulation of mRNA catabolic process, post-transcriptional regulation of gene expression, and DNA destabilization provides insight into the regulatory mechanisms that govern gene expression in response to FAE exposure in females. Cellular components included the actin cytoskeleton, preribosome, large subunit precursor, focal adhesion, and cell-substrate junction, which reflects alterations in cytoskeletal dynamics, cell adhesion and signaling pathways, and ribosome biogenesis, which may result in morphological changes. Molecular functions include RNA binding, DNA binding, telomerase activity, and mismatched DNA binding, which relates to DNA damage responses. The downregulated genes identified in the control female vs. FAE female analysis were associated with biological processes such as ncRNA processing, rRNA metabolic process, and ribosome biogenesis, indicating significant disruptions in RNA metabolism and protein synthesis. Additionally, processes like translation, cytoplasmic translation, and macromolecule biosynthetic processes highlight the impact of FAE on the cellular machinery responsible for protein production, suggesting a potential reduction in overall cellular function and viability in response to FAE exposure in females. Cellular components included the nucleolus, ribosome, and polysomal ribosome, reflecting alterations in ribosomal structure and function, which may lead to impaired protein synthesis. Molecular functions associated with the downregulated DEGs included rRNA binding, RNA binding, and NADH dehydrogenase activity, which relate to the cellular response to oxidative stress and energy metabolism. The downregulation of these functions suggests a compromised ability to manage oxidative damage and maintain homeostasis. Compared to the sex combined analysis, the female-specific analysis showed a more pronounced impact on ribosomal biogenesis and RNA-related processes. This suggests that while both analyses indicate disruptions in metabolic pathways due to FAE, the effects may manifest differently between sexes. Overall, these results suggest that FAE significantly affects ribosomal pathways, with strongest impacts observed in females.

### RNA-seq WGCNA confirms sex-specific dysregulations of gene modules associated with neurological activity

In addition to the DEG analysis, we further explored the relationships between gene expression patterns and sample traits using WGCNA. The subsequent analysis revealed distinct modules of co-expressed genes (Supplementary Table 4), which were correlated with the sample traits, including the pairwise comparisons derived from the differential expression analysis (Figure 4). We focused on modules that showed significant associations with FAE but not litter or sex-only effects. The blue module is significantly positively correlated with the control vs. FAE comparison, including both sex-stratified control vs. FAE comparisons (i.e. control female vs FAE female and control male vs FAE male). The genes within the blue module are associated with morphine addiction, the hedgehog signaling pathway, glutamatergic synapse, gap junction, focal adhesion, ECM-receptor interaction, dilated cardiomyopathy (DCM), circadian entrainment, and calcium signaling pathway KEGG terms (Figure 5). These findings underscore the critical role of the gene network represented by the blue module as positively associated with the biological effects of FAE. The turquoise module is significant for the same traits as the blue module, but negatively correlated instead, with additional significant KEGG terms including spliceosome, non-alcoholic fatty liver disease, Huntington disease, DNA replication, basal transcription factors, autophagy, and Alzheimer disease. The terms Huntington disease, Alzheimer disease, and autophagy highlight a strong correlation with neurological pathways.

**Figure 4.**
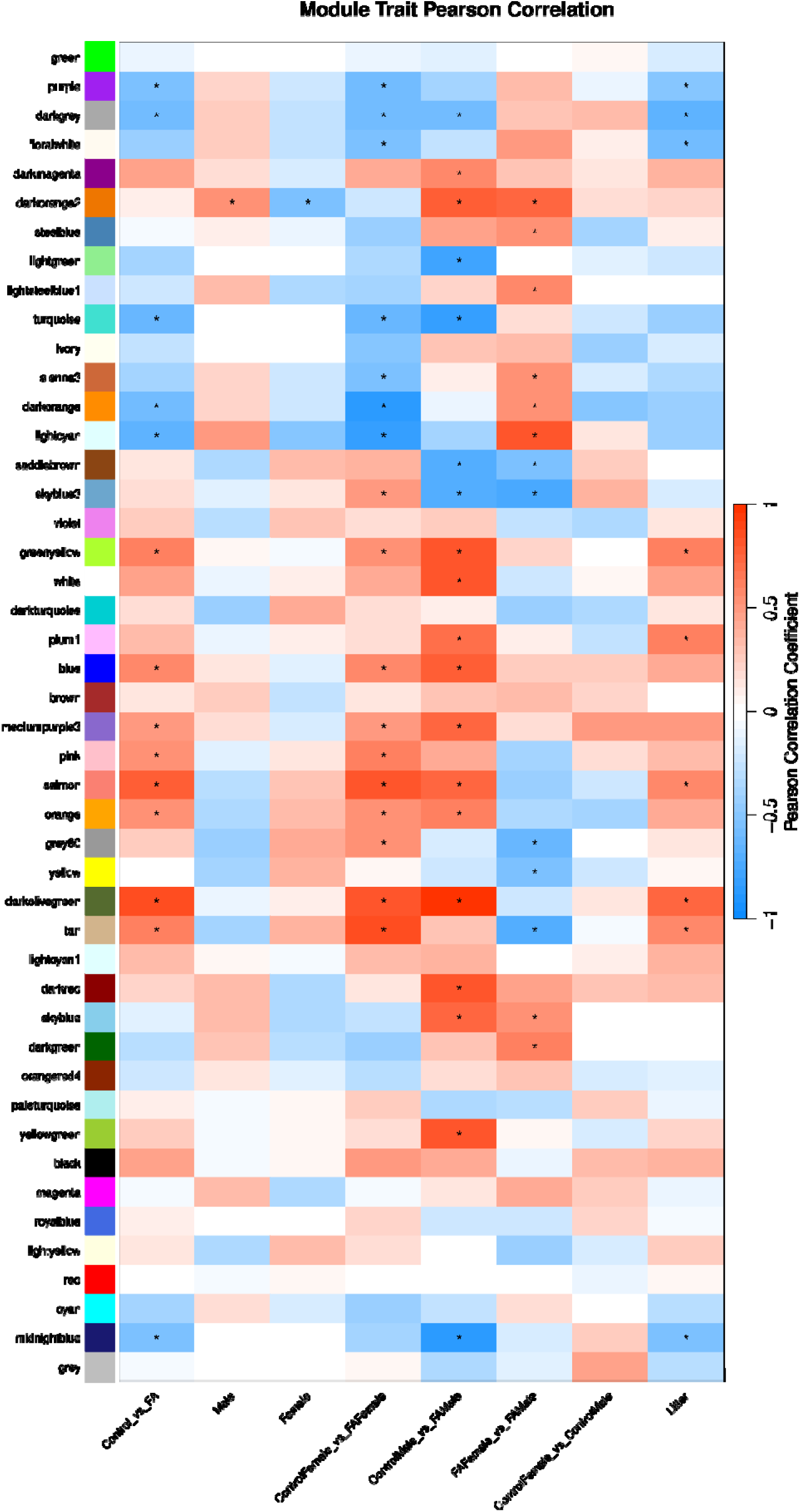
Heatmap of Module Trait Pearson Correlations from WGCNA. The heatmap displays the Pearson correlation coefficients between gene modules and various traits, including different experimental conditions and groups, sex, and litter. The color scale ranges from dark red, indicating a perfect positive correlation (1), to dark blue, indicating a perfect negative correlation (-1). Each cell in the heatmap represents the correlation coefficient for a specific module-trait pair, with darker shades of red signifying stronger positive correlations and darker shades of blue signifying stronger negative correlations. The traits analyzed include Control vs. FAE, Male, Female, Control Female vs. FAE Female, Control Male vs. FAE Male, FAE Female vs. FAE Male, Control Female vs. Control Male, and Litter. For all pairwise comparisons (e.g., Control vs. FAE, Control Female vs. FAE Female, Control Male vs. FAE Male), positive correlations (red) indicate that higher expression levels of specific gene modules are associated with the first group in the comparison. Conversely, negative correlations (blue) indicate that higher expression levels are associated with the second group. For the “Male” and “Female” traits, red indicates associations with male or female characteristics, respectively, while blue indicates associations with the opposite sex. In the “Litter” trait, red correlations suggest higher expression levels linked to specific litter groups, while blue indicates associations with other litter groups. Modules significant only in the context of litter were excluded from functional analysis. The stars represent significant correlations, highlighting relationships that are statistically meaningful. These significant correlations suggest that the associated gene modules have a notable impact on the traits being analyzed. This heatmap was made using R (v4.3.2) with an R kernel (r-irkernel v1.3.2) in a Jupyter Notebook (v7.0.7) with the following packages: tidyverse (v2.0.0), WGCNA (v1.72.5), magrittr (v2.0.3), and biomaRt (v2.58.2).

**Figure 5.**
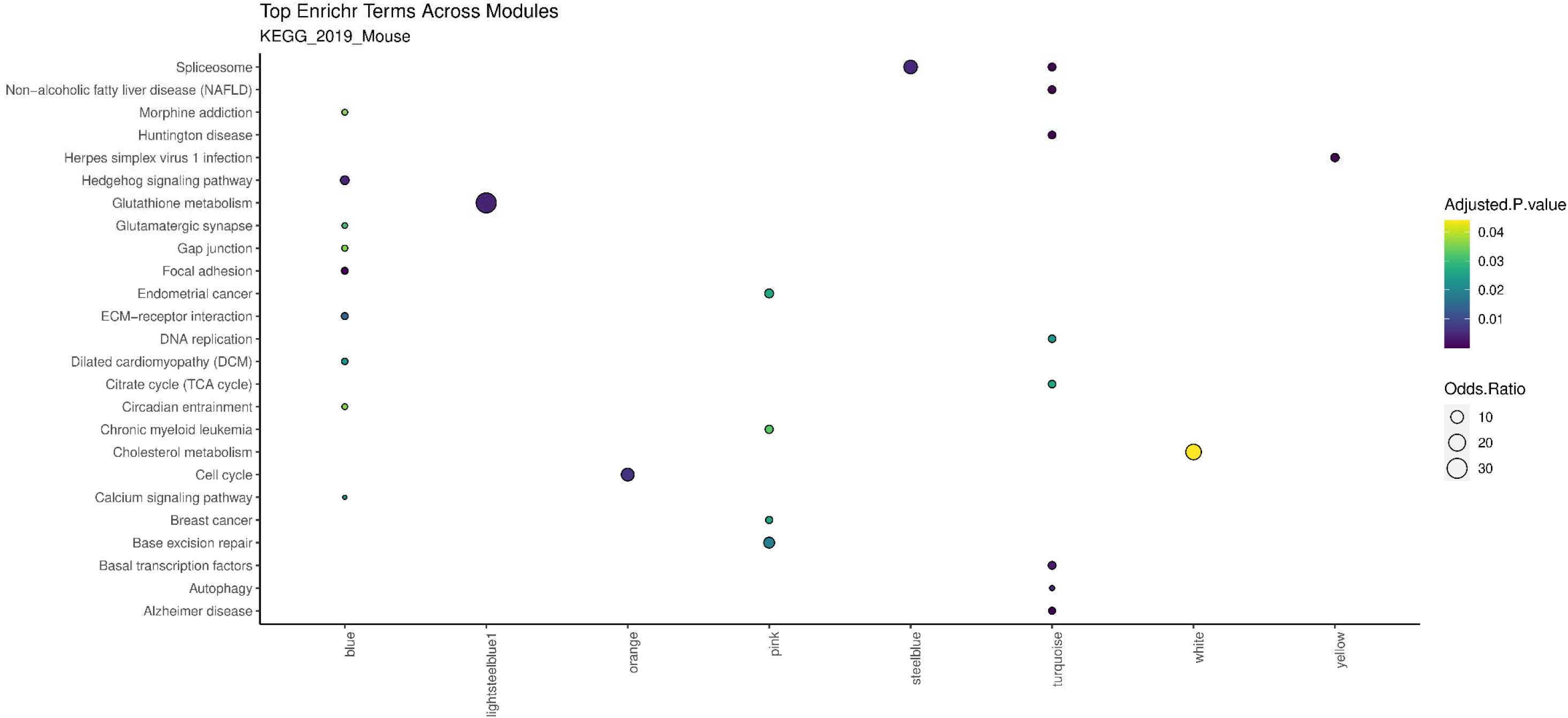
Dot Plot of Top KEGG Terms Across WGCNA Modules. This dot plot illustrates the top KEGG pathways identified across different gene modules derived from WGCNA. Each dot represents a specific KEGG term from the KEGG_2019_Mouse database being identified in the module. The size of the dot represents the odds ratio, with a larger dot (higher odds ratios) signifying a stronger association between the gene module and the KEGG pathway. Conversely, smaller dots (lower odds ratios) indicate weaker associations. The color of each dot reflects the adjusted p-value, with yellow indicating slight significance and darker blue shades indicating greater significance. The y-axis lists the various KEGG pathways, while the x-axis corresponds to the different WGCNA modules. Modules that were not significant for any condition or only for litter effects were excluded. This dot plot was made using R (v4.3.2) with an R kernel (r-irkernel v1.3.2) in a Jupyter Notebook (v7.0.7) with the following packages: readxl (v1.4.3), ggplot2 (v3.4.4), viridis (v0.6.5), dplyr (v1.1.4), glue (v1.7.0), and tidyr (v1.3.1).

The steelblue, lightsteelblue1, and darkgreen modules were also interesting due to their significant positive correlations only in the FAE female vs. FAE male comparison, meaning higher expression in FAE-exposed males than females. The steelblue gene network is enriched for genes associated with the spliceosome, and lightsteelblue1 was highly enriched for the glutathione pathway involved in antioxidant functions. Sex-specific associations with FAE were also seen in other modules, such as the pink module, that is significantly positively correlated with the control vs. FAE and control female vs. FAE female comparison, but not control male vs. FAE male. Pink module genes were enriched for genes associated with endometrial cancer, chronic myeloid leukemia, breast cancer, and base excision repair, suggesting that FAE may influence pathways related to cancer susceptibility and DNA repair pathways in females.

Overall, these results reveal significant insights into gene expression networks and the effects of FAE on these networks, highlighting both common and sex-specific pathways that may contribute to neurological and developmental differences.

### FAE alters DNA methylation levels in regions associated with neurodevelopmental processes and synapse activity

The WGBS analyses yielded 910 DMRs in the control vs. FAE comparison using sex as a covariate. The significant DMRs that covered 0.02% of the genome (Supplementary Table 5). 70% of the DMRs were hypermethylated, and 30% were hypomethylated. On average, DMRs were 576 base pairs long and contained 11 CpGs. DMRs were significantly enriched for exonic regions, with hypomethylated regions being both exonic and intronic (Figure 6A).

**Figure 6.**
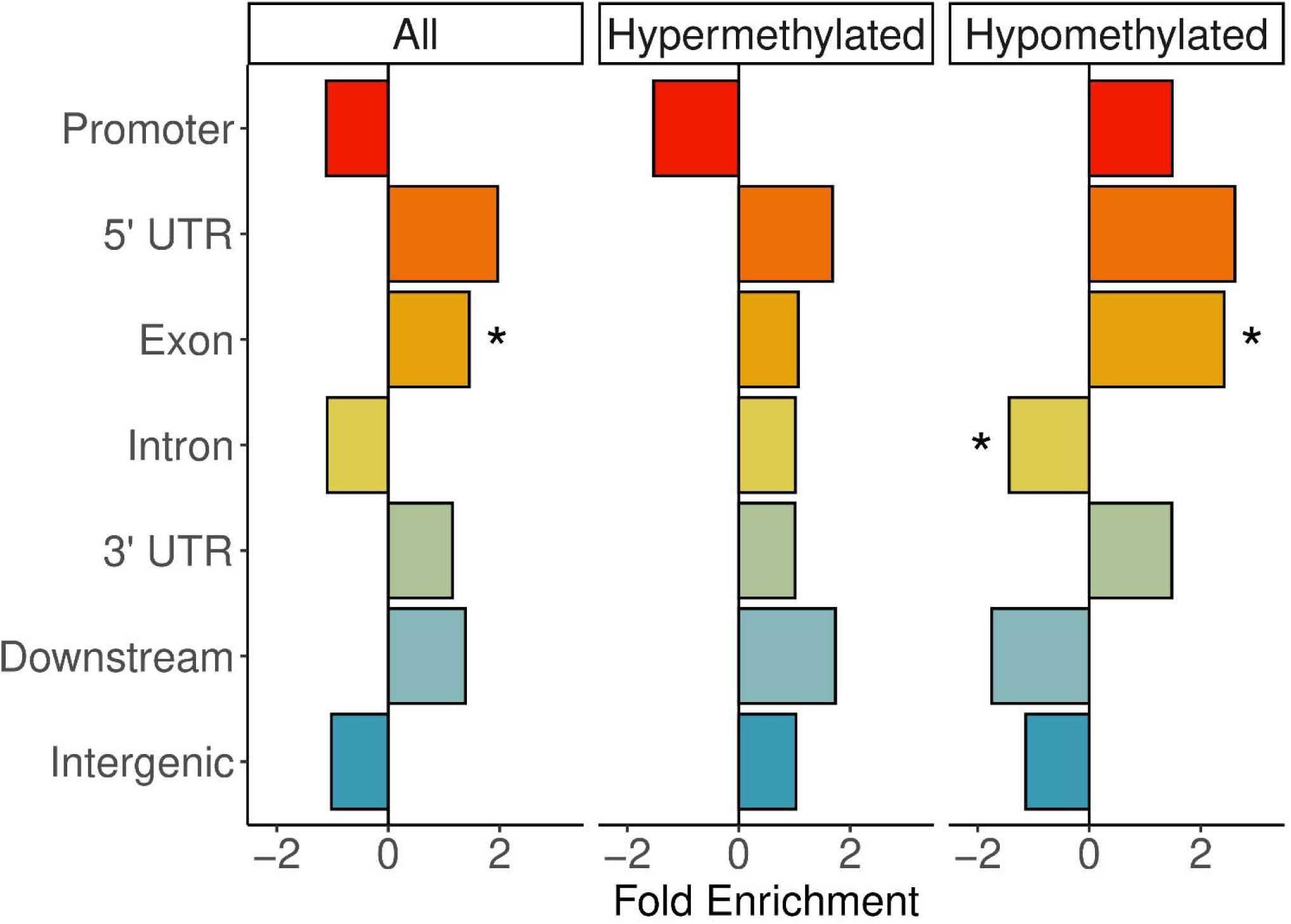
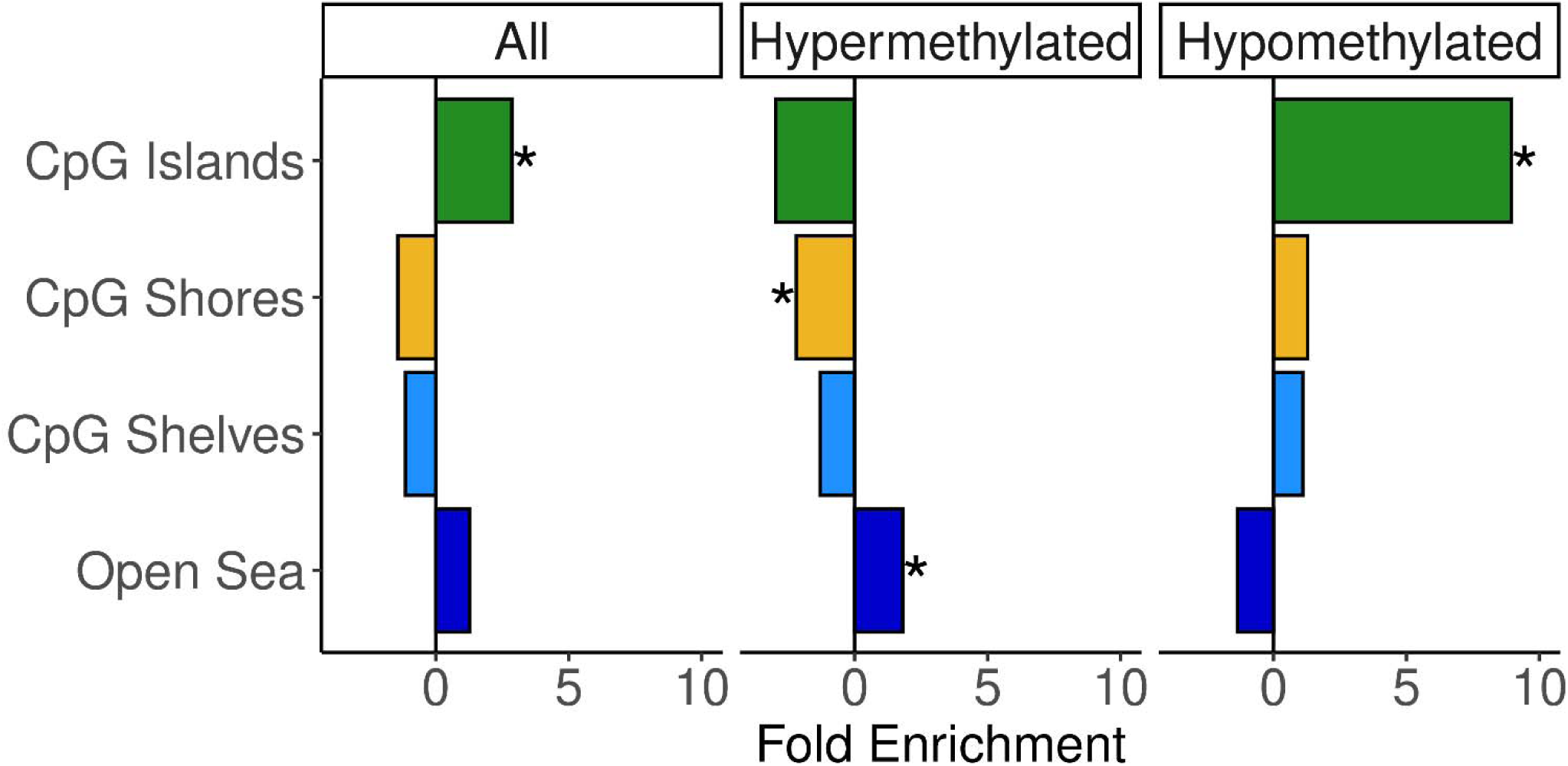
Genic Site and CpG Site Enrichment Based on DMRs. This figure presents an output from the DMRichR (v1.7.1) analysis, highlighting the enrichment of different genic and CpG sites based on the identified DMRs. **(A)** The x-axis represents the fold enrichment of hypermethylated and hypomethylated DMRs across various genic sites, including promoters (red), 5’ UTRs (orange), exons (orange-yellow), introns (yellow), 3’ UTRs (green), downstream regions (pale blue), and intergenic regions (blue). The y-axis categorizes the genic sites, with the bars indicating the degree of enrichment for hypermethylated and hypomethylated regions as well as a combination of all regions. Positive values on the x-axis (greater than 0) indicate a higher enrichment of DMRs in that specific genic site, while negative values (less than 0) suggest a depletion of DMRs. **(B)** While panel A focuses on genic sites, panel B illustrates the enrichment of DMRs across various CpG site categories, including CpG islands (green), CpG shores (yellow), CpG shelves (light blue), and open sea regions (dark blue). The x-axis represents the fold enrichment of hypermethylated and hypomethylated DMRs, while the y-axis categorizes the different CpG site types. Positive values indicate a higher enrichment of DMRs in that specific CpG category, while negative values indicate a depletion.

Hypermethylated DMRs were significantly enriched for CpG shores and the open sea and hypomethylated DMRs were significantly enriched for CpG islands (Figure 6B).

The top 10 DMRs with the highest methylation differences between the control and FAE groups were *Txndc2*, *Fstl4*, *Cmc1*, *Ubxn8*, *Gm35496*, *Cep250*, *Xpr1*, *Pitpnb*, *Fndc5*, *Adgrl2*, and *Shtn1*. *Txndc2* and *Cep250* are involved in oxidative stress responses and spermatogenesis, suggesting potential impacts on male fertility and reproductive health. *Fstl4* plays a role in cell differentiation and regulates BDNF, which is a critical player in neurodevelopment. *Cmc1* is associated with mitochondrial respiration, indicating that FAE may disrupt energy metabolism. *Ubxn8* is a ubiquitin regulatory protein that affects protein degradation, while *Xpr1* maintains phosphate homeostasis, both of which are important for metabolic regulation. *Pitpnb* is significant for phospholipid signaling in neurogenesis, and *Fndc5* is linked to insulin sensitivity. Lastly, *Adgrl2* and *Shtn1* are crucial for neuronal connectivity and function, particularly in neuronal polarization and axonogenesis. Together, these findings suggest that the epigenetic changes observed in these genes may contribute to adverse effects associated with FAE with regards to metabolic, cellular, and neurodevelopmental pathways.

Enrichment analysis of the DMRs identified significant associations with biological processes such as glutamate receptor signaling, axonogenesis, regulation of potassium ion transmembrane, positive regulation of neuron differentiation, and negative regulation of response to stimulus, indicating that FAE may disrupt critical pathways involved in neuronal development and function (Figure 7). Additionally, cellular components like the axon highlights the potential impact on neuronal structure. Molecular functions, including ionotropic glutamate receptor activity, RNA polymerase II transcription regulatory region activity, and neurotrophin binding suggest that epigenetic modifications resulting from FAE could alter gene expression and synaptic transmission. Collectively, these findings provide additional context on potential mechanisms by which FAE influences health outcomes, particularly in relation to neurological development and function.

**Figure 7.**
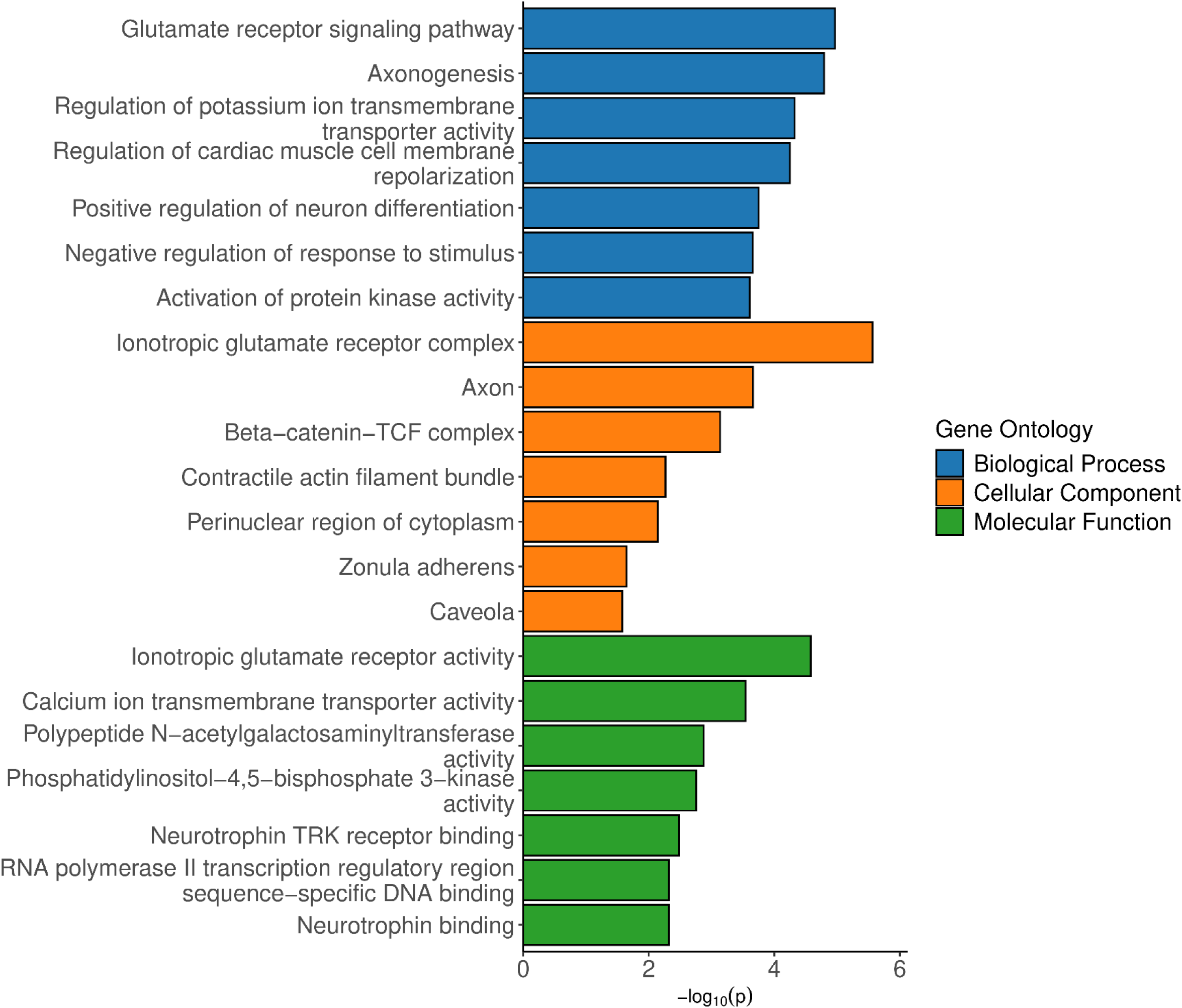
Enrichr Results for Gene Ontology Analysis of DMRs. Results of DMRichR (v1.7.1) analysis, specifically the gene ontology analysis conducted on significant DMRs. The y-axis lists the enriched GO terms categorized into three main categories: Biological Process (blue), Cellular Component (orange), and Molecular Function (green). The x-axis represents the significance of enrichment, indicated by the-log10(p-value), with higher values reflecting greater statistical significance.

### Comethylation region network analysis confirms dysregulated gene networks in FAE are predominantly associated with chemical synapse activity

Comparably to the WGCNA analysis for the RNA-seq data, we further explored the relationships between comethylated genes (i.e. modules) and sample traits using Comethyl.

This analysis yielded 58 modules, with the hondeydew1, white, brown, brown4, and purple modules being particularly interesting due to their significance in the FAE group (Figure 8, Supplementary Table 6). The brown, brown4, purple, and white modules were significantly enriched for KEGG terms related to neurological processes, including glutamatergic synapse, GABAergic synapse, dopaminergic synapse, cholinergic synapse, calcium signaling pathway, axon guidance, and more (Figure 9).

**Figure 8.**
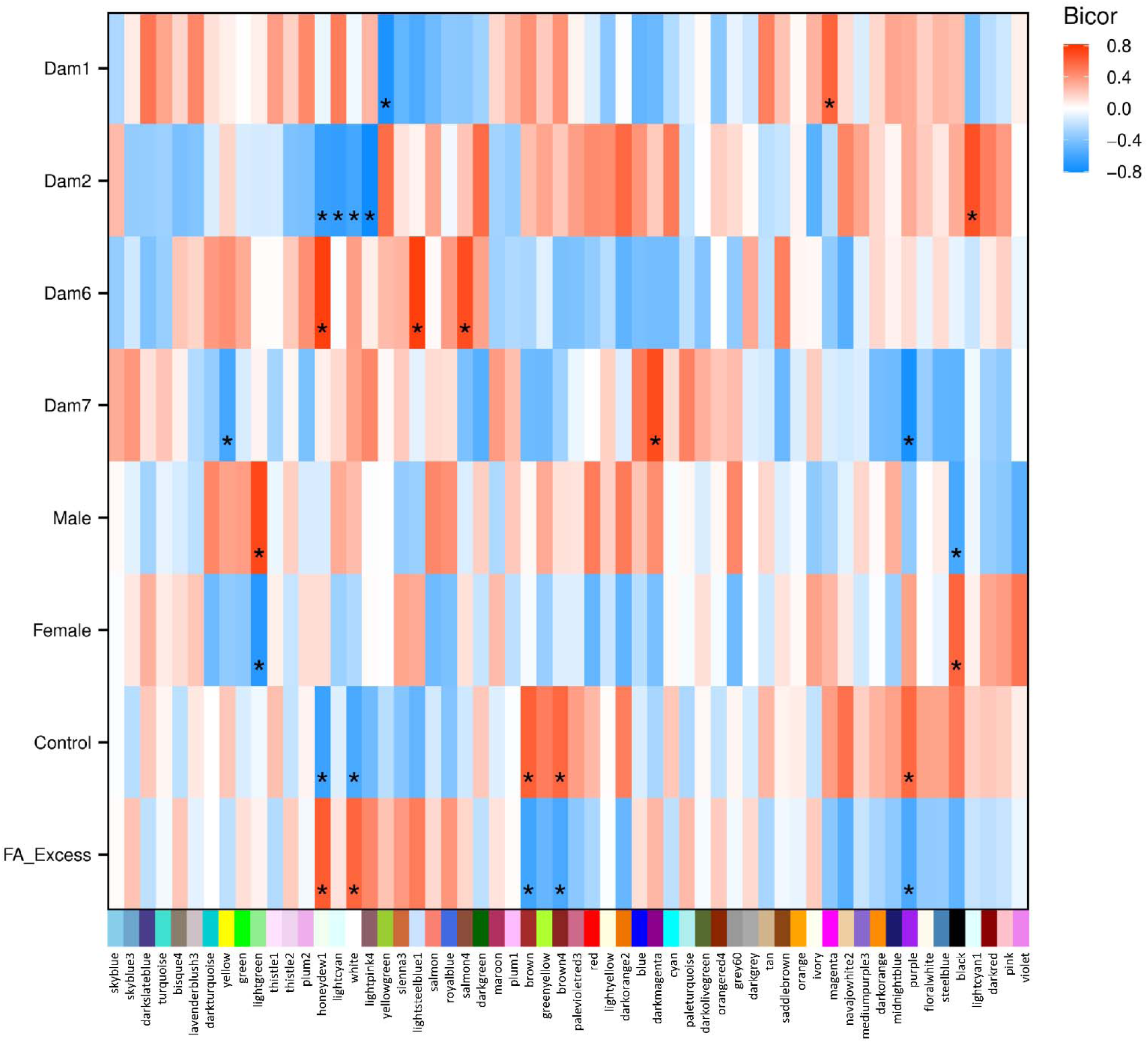
Heatmap of Module Trait Biweight Midvariance Correlations from Comethyl. This figure presents an output from the Comethyl (v1.3.0) analysis. This heatmap displays the biweight midvariance correlation coefficients between comethylated modules and various traits, including different the litter (“Dam”), sex, and experimental condition. The color scale ranges from red, indicating a perfect positive correlation (1), to blue, indicating a perfect negative correlation (-1). Each cell in the heatmap represents the correlation coefficient for a specific module-trait pair, with darker shades of red signifying stronger positive correlations and darker shades of blue signifying stronger negative correlations. The traits analyzed include Dam1, Dam2, Dam6, Dam7, Male, Female, Control, and Folic Acid Excess (“FA_Excess”). The stars represent significant correlations, highlighting relationships that are statistically meaningful. These significant correlations suggest that the associated comethylated modules have a notable impact on the traits being analyzed.

**Figure 9.**
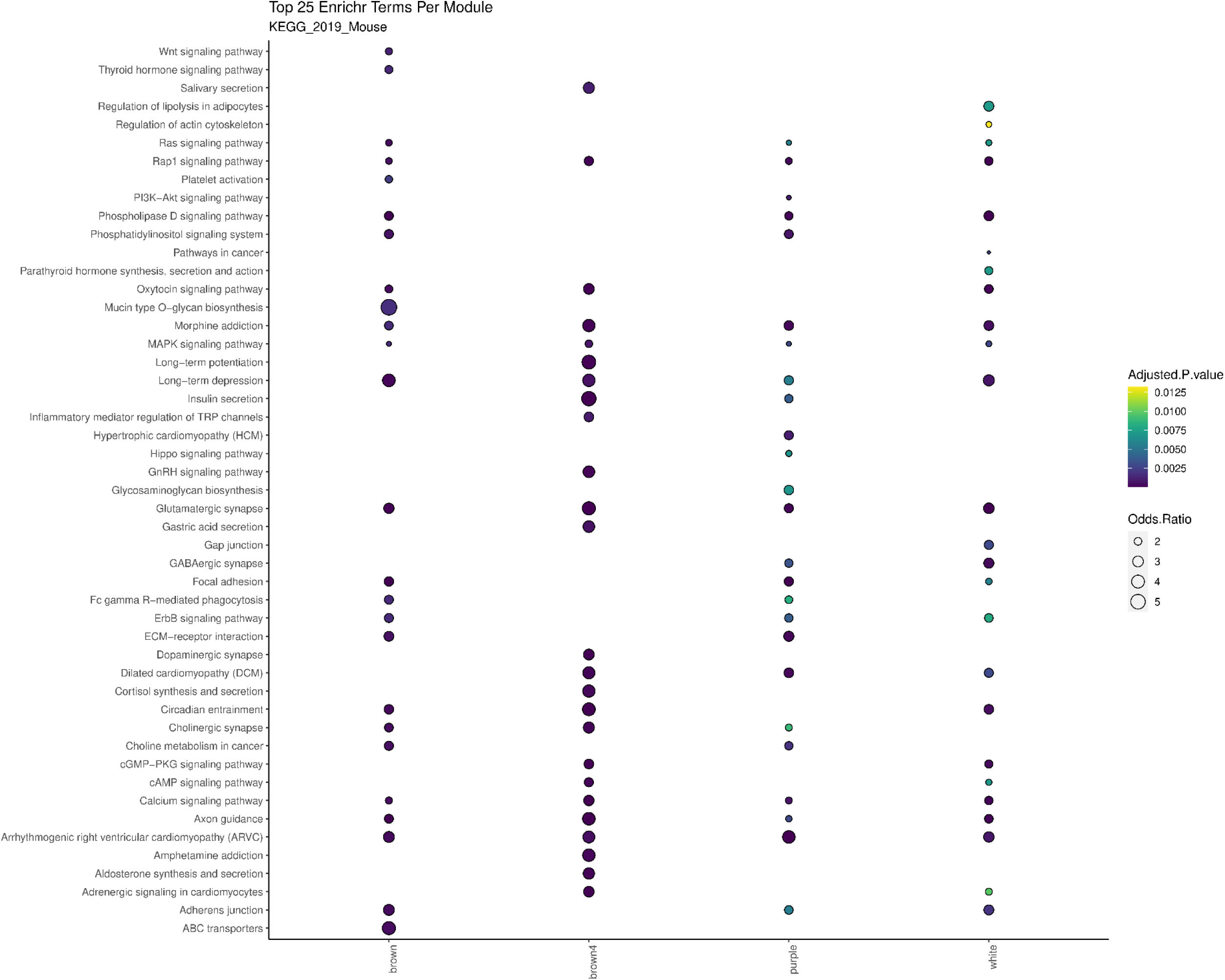
Dot Plot of Top KEGG Terms Across Comethyl Modules. This dot plot illustrates the top KEGG pathways identified across different gene modules derived from Comethyl. Each dot represents a specific KEGG term from the KEGG_2019_Mouse database being identified in the module. The size of the dot represents the odds ratio, with a larger dot (higher odds ratios) signifying a stronger association between the gene module and the KEGG pathway. Conversely, smaller dots (lower odds ratios) indicate weaker associations. The color of each dot reflects the adjusted p-value, with yellow indicating slight significance and darker blue shades indicating greater significance. The y-axis lists the various KEGG pathways, while the x-axis corresponds to the different Comethyl modules. Depicted modules were significant for the FAE condition containing significantly enriched KEGG terms. This dot plot was made using R (v4.3.2) with an R kernel (r-irkernel v1.3.2) in a Jupyter Notebook (v7.0.7) with the following packages: readxl (v1.4.3), ggplot2 (v3.4.4), viridis (v0.6.5), dplyr (v1.1.4), glue (v1.7.0), and tidyr (v1.3.1).

### Integration of RNA-seq DEGs and WGBS DMRs reveals significant correlations between data sets

Significant DEGs from the RNA-seq analysis were compared to the significant DMRs from the WGBS analysis for the control vs. FAE groups to identify overlaps. It is important to note that the RNA and DNA samples of each sequencing experiment were obtained from the same set of tissue samples. The integration of RNA-seq DEGs and WGBS DMRs revealed an overlap of 20 genes: *Naa20*, *Med10*, *Epb41l4a*, *Katnal2*, *D3Ertd751e*, *Ccdc93*, *Itga6*, *Tacr3*, *Cald1*, *Ccl17*, *Atp2c1*, *Asprv1*, *Kcnk10*, *Fau*, *Egr1*, *Mest*, *Fbh1*, *Syt10*, *Urm1*, and *Arrdc3*. Based on the Fisher’s Exact Test, this overlap is significant (p-value = 2.2 x 10^-16^). In addition to this overlap being significant, 809 genes also showed significant correlations between RNA-seq TPMs and WGBS percent methylation values (Supplementary Table 7). Among these significant transcriptome-methylome correlated genes, 583 were identified in control mice (553 uniquely, 238 positively correlated, 345 negatively correlated), compared to only 288 genes with transcriptome-methylome correlations in the FAE mice (256 unique to FAE group, 110 positively correlated, 178 negatively correlated). These results suggest that part of the dysregulation associated with prenatal FAE may be the disrupted balance of DNA methylation regulatory elements important for gene expression.

### FA imbalance disrupts network-level functional connectivity in mouse newborn cortical circuits

Our earlier work revealed that mice exposed prenatally to folate deficiency and FA excess show reduced dendritic arborization but increased synaptic density in cortical projection neurons^88,89^. Since dendrites and their spines are fundamental to neuronal connectivity and communication, these structural changes, in combination with findings here of altered gene networks relevant to chemical synapse function, may have consequences for neuronal physiology. To examine this question, we tested electrophysiological function using HD-MEAs on dissociated cortical cells derived from control and FAE newborn pups. HD-MEA recordings revealed marked differences in network activity between control and FAE-derived cortical neurons. Figure 10 shows representative raster plots from DIV21 recordings, illustrating distinct burst patterns differentiating both conditions. Control cultures exhibited robust, synchronized bursting patterns characteristic of healthy neuronal networks (Fenton et. al 2024) (Fig. 10a). In contrast, FAE-derived networks showed significant disruptions in synchronization, with lower burst amplitudes and altered burst frequencies (Fig. 10b). FAE neurons consistently displayed higher numbers of bursts coupled with substantially reduced burst peak amplitudes (Fig. 10c,d) compared to controls. These electrophysiological differences persisted throughout the recording period (DIV 7-33), indicating that prenatal exposure to excess FA induces persistent alterations in network-level neuronal communication. The observed reduction in burst amplitude and increased burst frequency in FAE cultures suggests compromised network synchronization at the cellular level^90,91^.

**Figure 10.**
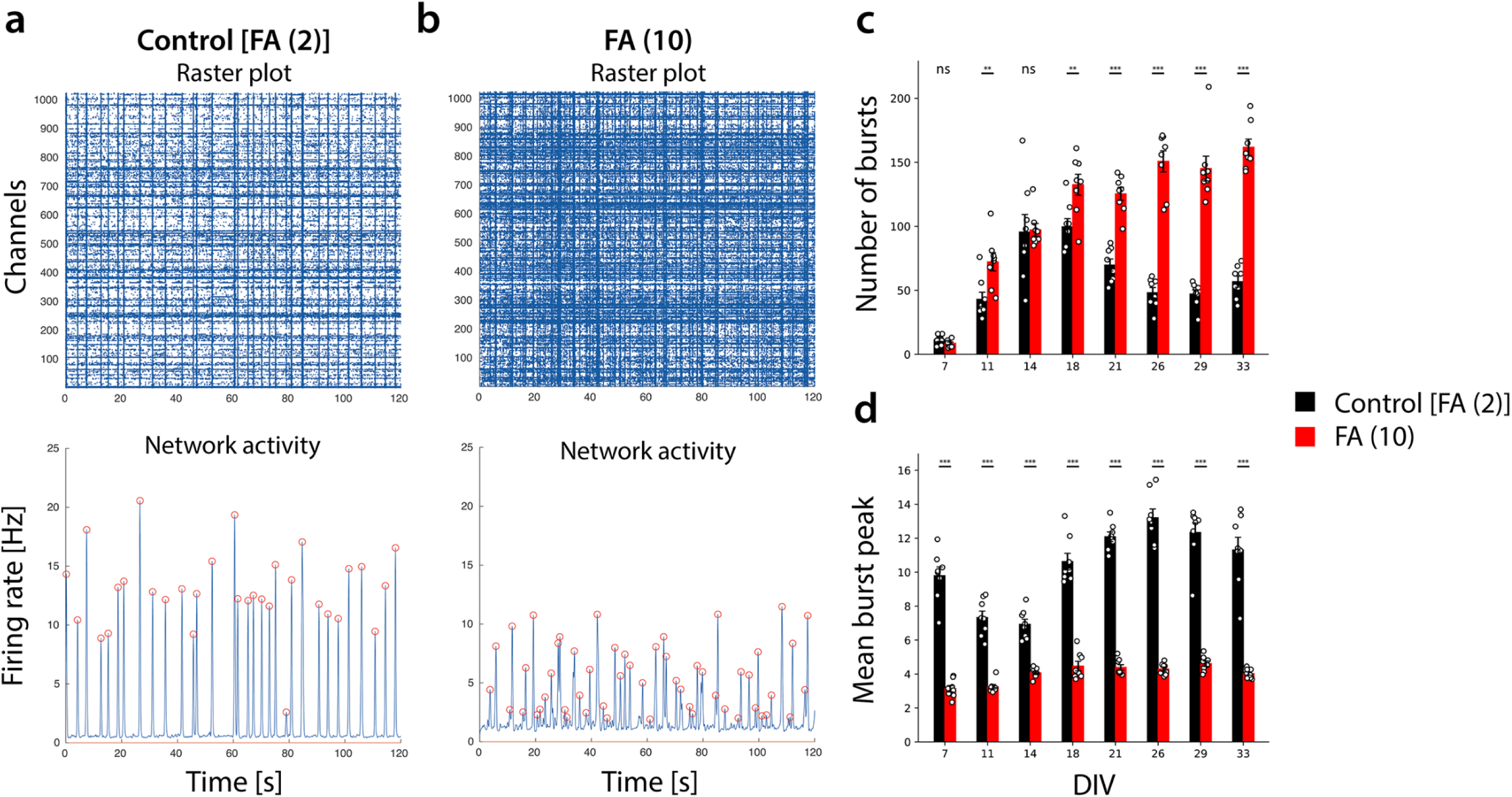
HD-MEA Network and Burst Activity. HD-MEA recordings reveal altered network activity in cortical neurons derived from FAE pups compared to controls. **(A)** Representative raster plots of network activity at DIV21 show distinct burst patterns over 120-second intervals, with FAE neurons (right) exhibiting reduced firing amplitude but increased burst frequency compared to controls (left). **(B)** Quantification of network burst frequency across development (DIV 7-33) demonstrates significantly increased burst frequency in FAE neurons (10 mg/kg, red) compared to controls (2 mg/kg, yellow). **(C)** Quantification of burst peak amplitudes shows substantial decreases in FAE neurons compared to controls throughout development.

## Discussion

The findings of this study provide insight into the biological consequences of 5-fold excess FA exposure during pregnancy on cerebral cortical DNA, methylation, gene expression, and network-level electrophysiology. Our analysis revealed significant alterations in gene networks associated with metabolic and neurological functions. The identification of 646 DEGs in the RNA-seq analysis highlights the impact of FAE on gene expression. Notably, the significant downregulation of genes critical for neurodevelopment, such as *Hhex, Lnp1* and *Dcst2*, and significant upregulation of *Ackr2*, raises concerns about the long-term implications of FAE on brain development and function. These genes are known to play pivotal roles in neuronal differentiation and the formation of neural circuits and synpases, suggesting that prenatal exposure to high levels of FA may disrupt normal neurodevelopmental trajectories. Furthermore, the RNA regulatory mechanisms identified in the female FAE GO analysis suggest that while we may not see significant changes at the RNA expression level, there could be substantial alterations at the proteomic level. These processes can lead to the degradation of specific mRNAs, preventing their translation into proteins, which may mask the true extent of gene expression changes when solely analyzing RNA levels. Additionally, indications of compromised genomic integrity may lead to aberrant protein synthesis or the production of dysfunctional proteins. This disconnect between RNA and protein expression highlights the need to investigate gene network level interactions and potentially expand into proteomic analysis methods to fully understand the biological consequences of FAE exposure, as the effects on protein function and interaction networks may be more pronounced.

The WGBS analysis identified 910 DMRs, with a predominance of hypermethylation, which is generally associated with gene silencing. The enrichment of DMRs in exonic regions and their association with critical biological processes, such as glutamate receptor signaling and axonogenesis, suggests that FAE may disrupt essential pathways involved in neuronal development and synaptic function. Furthermore, comethylated regions also suggested significance in several pathways involved in neurological function.

The molecular dysregulations uncovered by RNA-seq and WGBS provide also a possible explanation for the altered electrophysiological activity observed in FAE-derived neuronal networks. The electrophysiological deficits observed in FAE-derived cortical networks provide functional evidence of altered neuronal communication^92–94^. The reduced burst amplitudes and increased burst frequencies seen in FAE neurons reflect compromised network synchronization, which suggests that FAE exposure may disrupt fundamental mechanisms underlying neuronal circuit formation^95,96^. This pattern of altered burst dynamics - specifically, more frequent but weaker coordinated firing events - is consistent with impaired interneuronal communication and synchrony^97,98^. Similar network-level disruptions have been documented in other models of neurodevelopmental disorders^92,99,100^. The persistent nature of these electrophysiological abnormalities throughout in vitro development (DIV 7-33) indicates that prenatal FAE exposure induces long-lasting changes in neuronal network function that could contribute to altered neurodevelopmental trajectories.

Intriguingly, the reduced network synchronization and altered burst patterns, particularly reduced burst amplitude, observed in our HD-MEA recordings of FAE neurons closely resemble pattens observed previously in ASD. Particularly noteworthy is the alignment with seminal work by Bosl et al. who demonstrated decreased neural connectivity in infants at risk for ASD using EEG measurements^101^. While their study examined whole-brain connectivity patterns in human subjects and our investigation focused on cellular-level network formation in a mouse model, both approaches showed similar patterns of reduced neuronal synchronization and complexity. Our findings extend this observation suggesting that prenatal FAE can disrupt neuronal connectivity, both structurally by diminishing axonogenesis, and potentially through its impact on chemical synapse function and particularly its impact on glutamatergic synapses, one of the gene networks most severely impacted according to our analyses. Although our study, like the early EEG investigations, cannot definitively establish causality between altered connectivity and ASD, it provides mechanistic insights into how environmental factors such as FAE might influence early neurodevelopmental trajectories. The convergence of findings across different experimental scales - from molecular pathways to cellular networks - strengthens the hypothesis that disrupted neural connectivity during early development may represent a common pathway in neurodevelopmental disorders and particularly ASD.

The findings of this study also reveal important connections between FAE and blood-related health issues. The significant downregulation of *Lnp1*, a fusion partner of NUP98 associated with hematopoietic malignancies, underscores the potential cancer-related implications of FAE, aligning with results from other studies that have linked folate metabolism to increased cancer risk^102^. The increased cancer risk may be a result of enhanced cell growth and proliferation, as seen in the GO analysis for genes upregulated with FAE exposure. This is further supported by the Comethyl analysis, which identified significant enrichment for the following KEGG terms: adrenergic signaling in cardiomyocytes, arrhythmogenic right ventricular cardiomyopathy (ARVC), dilated cardiomyopathy (DCM), hypertrophic cardiomyopathy (HCM), and choline metabolism in cancer.

In addition to blood-related health outcomes, the presence of addiction-related GO terms in both the RNA-seq WGCNA and WGBS comethylation modules suggests that FAE may influence pathways relevant to substance use disorders. Current studies regarding the role of FA in addiction pathways suggest that FA deficiencies are an outcome of substance abuse^103–105^; however, it may be worth to further explore these associations to determine the impact of FAE and deficiencies on substance use disorders.

While the role of FA metabolism in blood-related health outcomes and addiction pathways warrants further investigation, previous studies have linked disruptions to insulin metabolism to FAE^41–43^. This aligns with our findings, which revealed significant hypomethylation of *Fndc5*.

*Fndc5* plays roles in the modulation of energy expenditure and insulin sensitivity. Hypomethylation is typically associated with increased gene expression, suggesting that the excessive expression of *Fndc5* may lead to dysregulation of insulin sensitivity pathways, contributing to insulin resistance. Thus, the hypomethylation of *Fndc5* provides a mechanistic insight into how altered folate metabolism may influence insulin sensitivity and overall metabolic health. In addition to the gene body level, comethylation modules significant for the FAE group had significant enrichment of genes involved in insulin secretion.

Interestingly, our results also revealed sex-specific differences in the response to FAE, with distinct patterns of gene expression and methylation observed. The identification of WGCNA modules that were positively correlated with FAE in sex-specific contexts suggests that the effects of FAE may manifest differently based on sex. This suggests that prenatal exposures can have differential impacts on male and female neurodevelopment, potentially contributing to the observed sex disparities in neurodevelopmental disorders such as ASD. The hypomethylation of *Txndc2* and *Cep250*, both linked to oxidative stress and spermatogenesis, supports a potential impact of prenatal FAE exposure on neurodevelopmental outcomes of male offspring. Conceivably, male offspring may be predisposed to adverse neurodevelopmental outcomes by interference with oxidative stress pathways that are critical for normal brain development^106^. Additionally, the alterations in gene expression and methylation patterns observed in this study suggest that FAE could negatively affect male reproductive health, potentially leading to fertility issues later in life. Understanding these sex-specific responses to FAE is vital for developing targeted nutritional guidelines and interventions that consider the unique metabolic and developmental needs of males and females.

Overall, our study provides novel insights into the molecular consequences of prenatal FAE exposure, revealing significant disruptions in gene expression and epigenetic regulation in the neonatal mouse brain. Further, our findings demonstrate that neurons derived from offspring exposed to FAE during pregnancy exhibit diminished capacity for establishing functional synaptic connections and creating coherent neuronal networks compared with controls. The integration of electrophysiological and molecular analyses provides compelling evidence that excess FA exposure during development may disrupt the fundamental mechanisms underlying neuronal communication and circuit formation. These findings highlight the importance of understanding the potential risks associated with high levels of FA intake and call for further research to determine the mechanisms underlying these effects. Future studies should aim to explore the long-term implications of these molecular changes and their relevance to the development of neurodevelopmental disorders, ultimately informing guidelines for FA supplementation in pregnant populations. The implications of these findings are significant, as they suggest that while FA fortification has been a successful public health measure in reducing neural tube defects, the potential risks associated with excessive intake after the periconceptional period of demonstrated protection warrant further investigation. The observed alterations in gene expression and methylation patterns may have long-term consequences for metabolic health, immune function, and neurodevelopment, underscoring the need for a balanced approach to FA supplementation during pregnancy.

## Conclusion

This study investigated the impact of prenatal FAE exposure on gene expression, DNA methylation, and electrophysiology patterns in the neonatal mouse brain. Our results demonstrate that FAE can lead to widespread alterations in gene networks critical for neurodevelopment, with potential long-term implications for metabolic health and neurological function, further illustrated by abnormal burst patterns of FAE-derived neuronal networks. The integration of RNA-seq and WGBS analyses revealed a complex interplay between epigenetic modifications and gene regulation, highlighting the interconnectedness of these processes in shaping developmental outcomes. Additionally, the identification of sex-specific differences in response to excessive prenatal FA exposure further underscores the necessity of considering offspring sex and timing of supplementation in research design and public health recommendations. While FA fortification and periconceptional use of prenatal vitamins has been instrumental in reducing neural tube defects, our results raise important questions regarding the potential risks associated with FAE in other aspects of neurodevelopment. In summary, this study contributes to the growing body of evidence suggesting that while FA is essential for healthy prenatal development, careful consideration of dosage and timing of administration is crucial to avoid unintended adverse effects, warranting further research to explore the long-term consequences of these molecular changes and to refine guidelines for FA supplementation in pregnant individuals, ensuring optimal health outcomes for both mothers and their offspring.

## Data Availability

Sequence data generated for this study can be found using the BioProject Accession Number PRJNA1223393. All scripts used in the RNA-seq and WGBS analysis can be found here: https://github.com/vhaghani26/Mouse_FAE_RNAseq_WGBS.

## Funding

This study was supported by NICHD R01HD107489, by a UC Davis Interdisciplinary Research Grant, and the Powell Family Charitable Trust. Additional support to KSZ was provided by Shriners Hospitals for Children and NIMH R21MH115347. Sara M. Ali was supported by a Ph.D. scholarship from the Egyptian Ministry of Higher Education and Scientific Research.

## Acknowledgements

The authors would like to thank Dr. Osman Sharifi, Dr. Aron Mendiola, and Dr. Blythe Durbin-Johnson for their insights on the RNA-seq and WGCNA analysis and Dr. Ian Korf for his computational insight and advice.

## Author Contributions

KSZ, JL, RG, and RBS devised and supervised this study. SMA and NC performed experiments. VH and SMA collected and analyzed data. VH, SMA, JL, RG, RBS, and KSZ co-wrote the manuscript.

## Conflict-of-Interest Declaration

The authors declare no conflicts of interest.

---

**Supplementary Table 1 – DEG Results for Control vs. FAE.** This table lists the DEGs identified in the RNA-seq analysis for the combined control vs. FAE analysis using sex as a covariate, including their Ensemble and common gene names (Ensemble Gene Name, Gene Name), log fold changes (logFC, positive is upregulated in FAE and negative is downregulated in FAE), average expression levels (AveExpr), t-statistics (t), p-values (P.Value), adjusted p-values (adj.P.Val, adjusted using the Benjamini-Hochberg method), B-statistics (B), and Entrez Gene IDs. Contrasts were conducted as groupFAexcess – groupControl.

**Supplementary Table 2 – DEG Results for Control Female vs. FAE Female.** This table lists the DEGs identified in the RNA-seq analysis for the control female vs. FAE female analysis, including their Ensemble and common gene names (Ensemble Gene Name, Gene Name), log fold changes (logFC, positive is upregulated in FAE females and negative is downregulated in FAE females), average expression levels (AveExpr), t-statistics (t), p-values (P.Value), adjusted p-values (adj.P.Val, adjusted using the Benjamini-Hochberg method), B-statistics (B), and Entrez Gene IDs. Contrasts were conducted as groupFAexcess.Female – groupControl.Female.

**Supplementary Table 3 – DEG Overlaps Between Groups.** This table presents the gene lists derived from the UpSet plot, categorizing DEGs based on various comparisons. The columns are organized to separately list significant downregulated and upregulated DEGs for the following comparisons: Control Female vs. FAE Female, Control vs. FAE, and the intersections of these comparisons.

**Supplementary Table 4 – Gene Composition of WGCNA Modules.** This table lists the genes associated with each WGCNA module, with each column representing a different module (i.e. genes in the same column belong to the same module).

**Supplementary Table 5 – DMR Results.** This table presents the full list of DMRs output by DMRichR (v1.7.1). Each row corresponds to a specific DMR and includes columns of associated data. The start and end columns indicate the genomic positions of the DMR, while the width column represents the length of the DMR, calculated as the difference between the end and start positions. The CpGs column specifies the number of CpG sites within the DMR. The betaCoefficient reflects the estimated change in methylation level, with positive values indicating hypermethylation and negative values indicating hypomethylation. The statistic column provides the statistical value associated with the significance of the methylation change, and the p.value indicates the statistical significance of the DMR, with lower values suggesting stronger evidence against the null hypothesis of no difference in methylation. The q.value is the adjusted p-value that accounts for multiple testing, offering a measure of significance that controls for false discovery rate. The direction column indicates whether the DMR is hypermethylated or hypomethylated in the FAE group relative to the control group, while the difference column shows the difference in methylation levels between the groups being compared. The absolute.value.difference provides the magnitude of change regardless of direction. The columns for CpG.Island, CpG.Shore, CpG.Shelf, and Open.Sea indicate whether the DMR is located within these specific genomic contexts. The annotation column describes the genomic context of the DMR (e.g., Exon, Intron, Distal Intergenic, 3’ UTR, etc.), and the geneId is the unique identifier for the gene associated with the DMR. The distanceToTSS column shows the distance from the DMR to the transcription start site (TSS) of the nearest gene, with negative values indicating that the DMR is upstream of the TSS. The ENSEMBL column provides the ENSEMBL gene identifier for the associated gene, while the geneSymbol and gene columns present the official gene symbol and the full name of the associated gene, respectively.

**Supplementary Table 6 – Gene Composition of Comethyl Modules.** This table lists the genes associated with each Comethyl module, with each column representing a different module (i.e. genes in the same column belong to the same module).

**Supplementary Table 7 – Spearman Coefficients for RNA-seq TPMs and WGBS Percent Methylation Per Gene.** The table presents the Spearman correlation coefficients and associated p-values for both control and FAE conditions. The “Gene” column lists the gene name. The “Spearman Correlation Coefficient for FAE” quantifies the strength and direction of the relationship between gene expression (measured in TPM) and percent methylation levels in the FAE condition, with positive values indicating a direct relationship and negative values indicating an inverse relationship. Coefficients close to 1 or-1 suggest strong correlations, whereas coefficients near 0 indicate weak or no correlation. The “FAE Correlation P-Value” indicates the statistical significance of the observed correlation for the FAE condition, where a lower p-value suggests a more significant correlation. The “Spearman Correlation Coefficient for Control” reflects the relationship between gene expression and percent methylation levels in the control mice, with similar interpretation as the FAE coefficient. The “Control Correlation P-Value” indicates the statistical significance of the correlation observed in the control group. The “DMR Distance to TSS” provides the distance of the DMR from the transcriptional start site of the gene. The “DMR Region Annotation” offers a description or classification of the DMR region, providing context about its genomic location. “Methylation Direction” indicates the direction of methylation change (e.g., hypermethylation or hypomethylation) associated with the DMR. The “RNA-seq Log Fold Change” indicates the change in gene expression between conditions where positive values are upregulated in FAE and negative values are downregulated in FAE. The “RNA-seq Adjusted P-Value” reflects the statistical significance of the log fold.

## Notes

https://github.com/vhaghani26/Mouse_FAE_RNAseq_WGBS

### Competing Interest Statement

The authors have declared no competing interest.

